# *In vivo* photopharmacology with light-activated opioid drugs

**DOI:** 10.1101/2023.02.02.526901

**Authors:** Shannan P. McClain, Xiang Ma, Desiree A. Johnson, Caroline A. Johnson, Aryanna E. Layden, Jean C. Yung, Susan T. Lubejko, Giulia Livrizzi, X. Jenny He, Jingjing Zhou, Emilya Ventriglia, Arianna Rizzo, Marjorie Levinstein, Juan L. Gomez, Jordi Bonaventura, Michael Michaelides, Matthew R. Banghart

## Abstract

Traditional methods for site-specific drug delivery in the brain are slow, invasive, and difficult to interface with recordings of neural activity. Here, we demonstrate the feasibility and experimental advantages of *in vivo* photopharmacology using “caged” opioid drugs that are activated in the brain with light after systemic administration in an inactive form. To enable bidirectional manipulations of endogenous opioid receptors *in vivo*, we developed PhOX and PhNX, photoactivatable variants of the mu opioid receptor agonist oxymorphone and the antagonist naloxone. Photoactivation of PhOX in multiple brain areas produced local changes in receptor occupancy, brain metabolic activity, neuronal calcium activity, neurochemical signaling, and multiple pain- and reward-related behaviors. Combining PhOX photoactivation with optical recording of extracellular dopamine revealed adaptations in the opioid sensitivity of mesolimbic dopamine circuitry during chronic morphine administration. This work establishes a general experimental framework for using *in vivo* photopharmacology to study the neural basis of drug action.

**Highlights:** A photoactivatable opioid agonist (PhOX) and antagonist (PhNX) for *in vivo* photopharmacology.

Systemic pro-drug delivery followed by local photoactivation in the brain.

*In vivo* photopharmacology produces behavioral changes within seconds of photostimulation.

*In vivo* photopharmacology enables all-optical pharmacology and physiology.

## Introduction

Pharmacological probes are widely used to study the nervous system. However, due to diffusion, traditional small molecule drugs act slowly and with spatial imprecision, even when applied locally in the brain through implanted cannulas. This impedes neuropharmacological studies *in vivo*, particularly those involving electrophysiological, imaging, and behavioral tracking measurements at high temporal resolution. Such studies greatly benefit from the ability to correlate measurements with well defined, time-locked stimuli that can be readily varied in intensity and duration. Photopharmacological reagents, which are converted by light from an inactive, pro-drug form into a biologically active form, potentially offer a general solution to this problem (Ellis-Davies, 2007; Velema, Szymanski and Feringa, 2014; Broichhagen, Frank and Trauner, 2015). As has been made clear by the widespread use of optogenetics, the use of light to control biological signaling provides experimental advantages due to the spatiotemporal precision of illumination and the ability to target deep brain structures in rodents using minimally invasive implanted optical fibers. The spatiotemporal precision of photopharmacology arises from the ability to pre-equilibrate brain tissue with the inactive drug form, such that illumination with brief (ms-sec) light flashes releases the active drug directly to its site of action, with sub-second kinetics, and in a highly stereotyped manner. Importantly, the local dose of active drug can be readily controlled by varying the amount of pro-drug administered or the amount of light delivered. These features of *in vivo* drug photoactivation have the potential to streamline studies involving local drug administration, especially in the context of ongoing animal behavior. Indeed, the first example of *in vivo* optogenetic control of animal behavior involved photoactivation of a caged ligand for a foreign receptor (Lima and Miesenböck, 2005). Furthermore, photopharmacology enables conceptually novel experiments built around the availability of a time-locked pharmacological stimulus that can be precisely controlled by the experimenter. However, despite a sizable pharmacopeia of photopharmacological probes constructed from molecular photoswitches and photochemical “caging” groups, many of which have been shown to work *in vivo* (Levitz *et al*., 2016; Font *et al*., 2017; Hüll, Morstein and Trauner, 2018; Acosta-Ruiz *et al*., 2020; Gomila *et al*., 2020; Donthamsetti *et al*., 2021), their potential as tools to study biological mechanism *in vivo* has yet to be fully realized. This shortcoming stems from limitations in pro-drug bioavailability, residual activity of the inactive form, and the requirement for long illumination periods due to poor bioavailability and/or inefficient photoactivation.

An attractive target for the development of photopharmacological probes that bypass these limitations is the mu opioid receptor (MOR), which is the primary target of morphine and other addictive analgesic drugs. In addition to MORs being important drug targets, endogenous opioid neuropeptides are implicated in diverse functions including feeding, memory, behavioral reinforcement, social interactions, and pain modulation. In particular, the neural mechanisms by which MOR agonists suppress pain and produce behavioral reinforcement are areas of intense investigation, within the overarching goal to develop non-addictive treatments for pain. One major area of focus is to understand the neural circuit adaptations to sustained drug use that underlie tolerance, dependence, withdrawal, and drug seeking behavior, which likely involves opioid-dopamine interactions. Yet, it has been challenging to evaluate drug-dependent changes in opioid-dopamine coupling within specific brain regions due to the imprecision and cumbersomeness of traditional methods for local drug delivery and measurement of dopamine signaling in awake, behaving animals. Our prior work with photopharmacological probes for MORs and other opioid receptors in *ex vivo* preparations (*e.g.* acute brain slices) has provided insights into mechanisms of neuropeptide polypharmacology, receptor interactions, signaling kinetics, and cellular adaptations to chronic opioid treatment (Banghart and Sabatini, 2012; Banghart *et al*., 2013; Williams, 2014; Banghart, He and Sabatini, 2018; He *et al*., 2021). However, these tools have yet to be applied *in vivo*.

Here, we describe a pair of photopharmacological reagents that afford spatiotemporally precise control of endogenous MORs *in vivo* using light. These molecules can be administered systemically and photoactivated with sub-second flashes of light through implanted optical fibers in the brain to drive rapid changes in behavior and neural circuit function. Importantly, because these reagents are compatible with commonly used *in vivo* optical methods, this work is an important step toward achieving all-optical manipulation and recording of neural function.

## Results

### Design, synthesis, and *in vitro* inactivity of PhOX and PhNX

To streamline our design and synthesis, we pursued photoactivatable analogues of the closely related agonist and antagonist oxymorphone (OXM, **1**) and naloxone (NLX, **2**), respectively (**Figure 1A**). We previously established that NLX can be readily caged via modification with a carboxynitroveratryl (CNV) group, yielding (CNV-NLX, **3**), which contains a negatively charged carboxylic acid at neutral pH (Banghart *et al*., 2013). To improve blood-brain barrier (BBB) penetrance, we exchanged the CNV group for a dimethoxynitrophenethyl (DMNPE) caging group (Wootton and Trentham, 1989), which contains a neutral methyl group in place of the carboxylic acid, affording photoactivatable oxymorphone (PhOX, **4**) and photoactivatable naloxone (PhNX, **5**). Although these caging groups require ultraviolet (UV) light for photoactivation, because they are insensitive to visible light, they are compatible with green and red fluorescent probes that are commonly used in *in vivo* imaging experiments (**Figure S1A**). Alkylation of OXM (**1**) with 1-(1-bromoethyl)-4,5-dimethoxy-2-nitrobenzene afforded PhOX (**4**) in 63% yield (**Figure 1B**). NLX (**2**) was similarly converted to PhNX (**5**) in 67% yield. Using high-pressure liquid chromatography and mass spectrometry, we confirmed that both PhOX and PhNX are stable in the dark for 24 hrs when dissolved in phosphate-buffered saline (PBS), and cleanly photoconvert back into OXM and NLX, respectively, upon exposure to 375 nm light (**Figure 1C**, **Figure S1B-D**). Similar to a recent study with a photo-activatable morphine analogue (López-Cano *et al*., 2021), we also prepared a derivative of OXM using the diethylaminocoumarin (DEAC) caging group (DEAC-OXM, **6**, **Figure 1A**). However, rather than cleanly converting to OXM upon UV illumination, DEAC-OXM underwent a 1,4 photo-Claisen rearrangement to predominantly yield a functionally-dead compound (RE-DEAC-OXM, **Figure S2, and Supporting Chemistry Data**), similar to other coumarin-caged phenols (Schaal *et al*., 2009; Wong *et al*., 2017). We therefore restricted further efforts to characterizing PhOX and PhNX.

**Figure 1.**
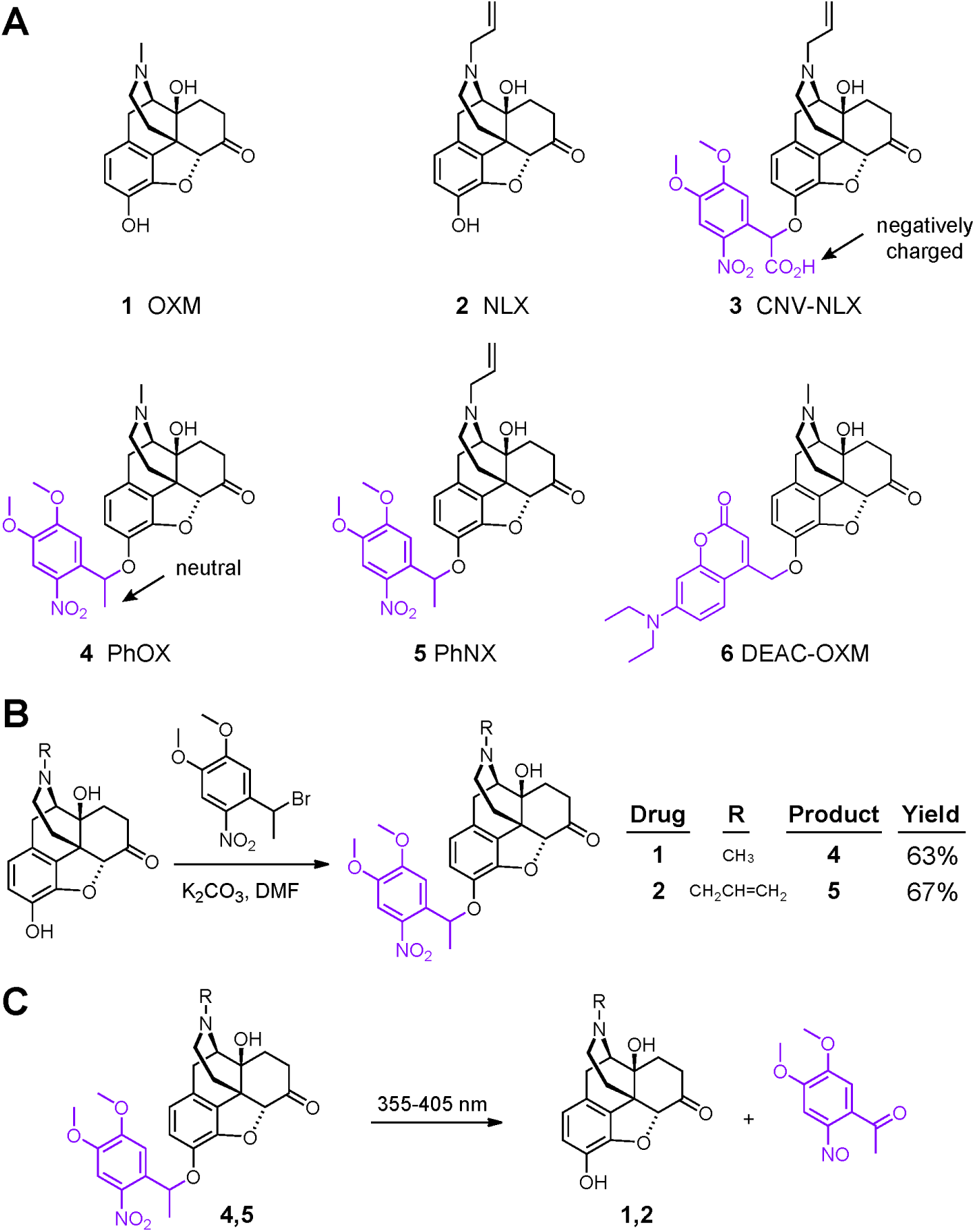
Design and synthesis of the photoactivatable oxymorphone and naloxone derivatives PhOX and PhNX. **(A)** Chemical structures of oxymorphone (OXM, **1**), naloxone (NLX, **2**), and the photoactivatable small molecule opioid drugs CNV-NLX (**3**), PhOX (**4**), PhNX (**5**), and DEAC-OXM (**6**). The light-removable The 1-(4,5-dimethoxy-2-nitrophenyl)ethyl (DMNPE) and 7-diethylaminocoumarinyl-4-methyl (DEAC) caging groups are drawn in violet. **(B)** Reaction scheme depicting the one-step alkylation procedure used to synthesize PhOX and PhNX from commercially available OXM and NLX, respectively. **(C)** Reaction scheme depicting ultraviolet light-driven photorelease of OXM and NLX from PhOX, and PhNX, respectively.

To determine if PhOX and PhNX exhibit reduced activity at MOR compared to OXM and NLX, we obtained dose-response curves at the murine MOR (SSF-MOR plasmid (Tanowitz and Von Zastrow, 2003)) expressed in human embryonic kidney cells, using the GloSensor™ assay, which reports cAMP level modulation by G protein signaling. Whereas OXM exhibited an EC50 of 1.4±0.2 nM and full receptor activation compared to the MOR agonist [D-Ala^2^, NMe-Phe^4^, Gly-ol^5^]enkephalin (DAMGO, 1 µM), at the highest concentration tested, PhOX (10 µM) produced only partial receptor activation, such that an EC50 was not determined (**Figure 2A**). Similar to CNV-NLX (Banghart *et al*., 2013), whereas NLX antagonized DAMGO (100 nM) with an IC50 of 86±28 nM, PhNX (up to 10 µM) exhibited no activity (**Figure 2B**). Dose-responses curves of DAMGO in the absence and presence of PhOX (300 nM), using NLX (100 nM) as a positive control, revealed that PhOX does not antagonize MOR at high but non-agonistic concentrations (**Figure 2C**). Further evaluation using a NanoBiT®-based luminescence complementation assay of β-arrestin signaling demonstrated that PhOX is not a biased agonist of the β-arrestin pathway (**Figure 2D**). Together, these results indicate that both PhOX and PhNX are highly inactive at MORs, yet cleanly provide OXM and NLX upon illumination with UV light.

**Figure 2.**
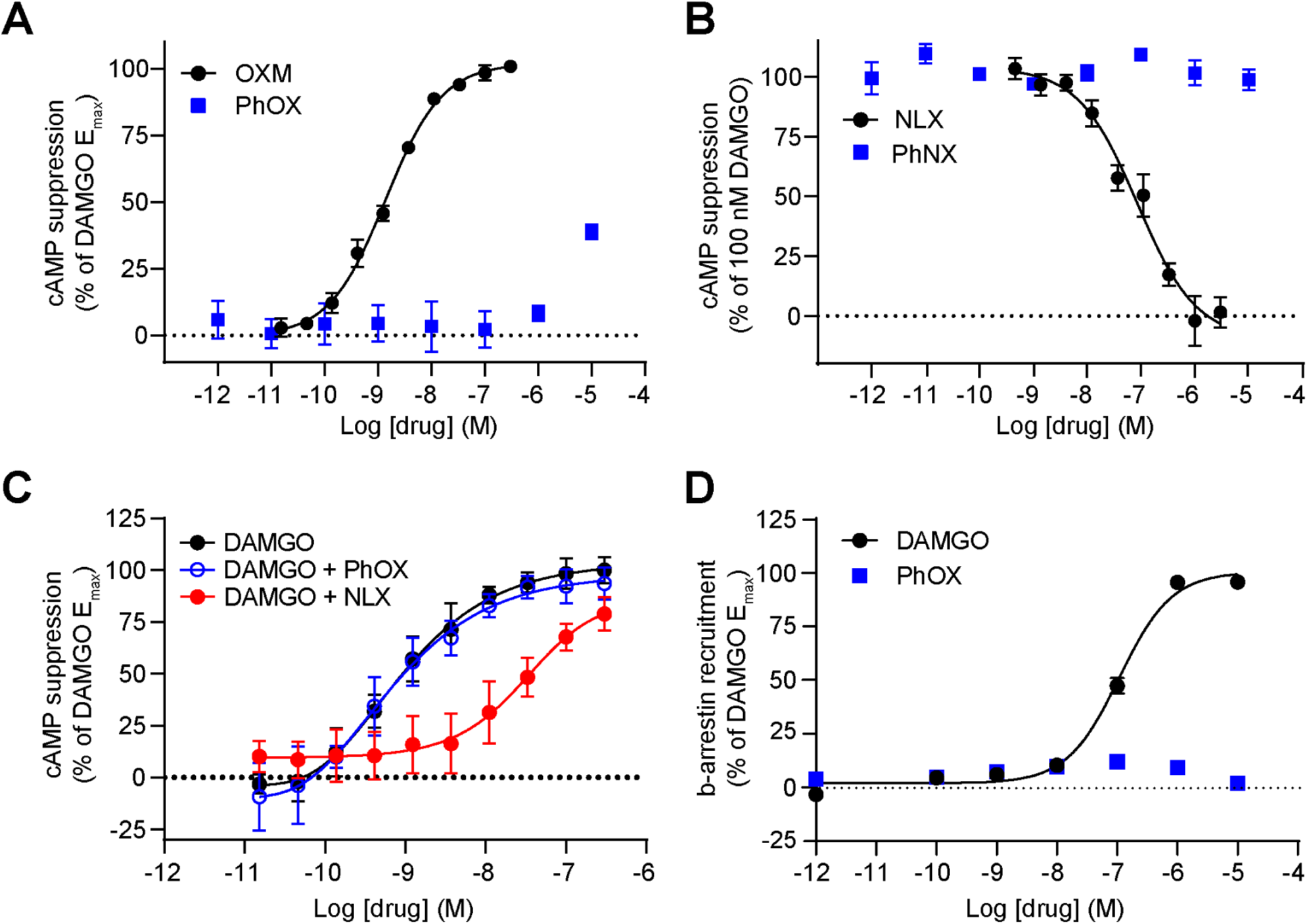
Evaluation of PhOX and PhNX residual activity at mu opioid receptors *in vitro*. **(A)** Agonist dose-response curves at the mu opioid receptor (MOR) for PhOX and OXM using the GloSensor™ assay of cAMP signaling in HEK293T cells. The solid line depicts the best-fit sigmoidal function used to derive the indicated EC_50_ value. OXM EC_50_ = 1.4 nM. Data were normalized to the response produced by DAMGO (1 µM) and are expressed as mean ± SEM (n=5 wells per concentration). **(B)** Antagonist dose-response curves at the MOR for NLX and PhNX in the presence of DAMGO (100 nM) using the GloSensor™ assay. Data are presented as in A. NLX IC_50_ = 86 nM. **(C)** Agonist dose-response curves at the MOR for DAMGO in the absence and presence of NLX (100 nM) or PhOX (300 nM) using the GloSensor™ assay. Data are presented as in A. DAMGO EC_50_ = 0.7 nM, DAMGO + PhOX EC_50_ = 0.9 nM, DAMGO + NLX EC_50_ = 32 nM. **(D)** Agonist dose-response curves at the MOR for DAMGO and PhOX using a NanoBiT-based luminescence complementation assay of β-arrestin signaling in HEK293T cells (n=4 wells per concentration). DAMGO EC_50_ = 11 nM. Data were normalized to the maximal response to DAMGO (10 µM) and are expressed as the mean ± SEM.

### Photoactivation of PhOX and PhNX engages endogenous opioid receptors *ex vivo*

We next asked whether photoactivation of PhOX and PhNX engages endogenous opioid receptors in acute brain slices taken from the CA1 region of mouse hippocampus, where MOR activation strongly suppresses perisomatic inhibition onto pyramidal neurons (Glickfeld, Atallah and Scanziani, 2008; He *et al*., 2021). During whole cell voltage clamp recordings, outward inhibitory post-synaptic potentials (IPSCs) were evoked by placing a small bipolar stimulating electrode adjacent to the recorded pyramidal cell, and isolated using antagonists for ionotropic glutamate receptors (**Figure 3A**). Whereas bath application of OXM (3 µM) suppressed IPSC amplitude by ∼60%, PhOX (3 µM) was inactive in the absence of light (**Figure 3B**). In the presence of PhOX (3 µM), application of a brief light flash from a 365 nm LED (50 ms, 5 mW) produced a rapid suppression of IPSC amplitude (∼40% suppression) (**Figure 3C**). This effect was blocked by NLX (3 µM), indicating the involvement of opioid receptors. These results are summarized in **Figure 3D**. Consistent with a pre-synaptic locus of opioid modulation, both the OXM- and PhOX-driven IPSC suppression were similarly associated with an increase in the paired pulse ratio (PPR, **Figure 3E**).

**Figure 3.**
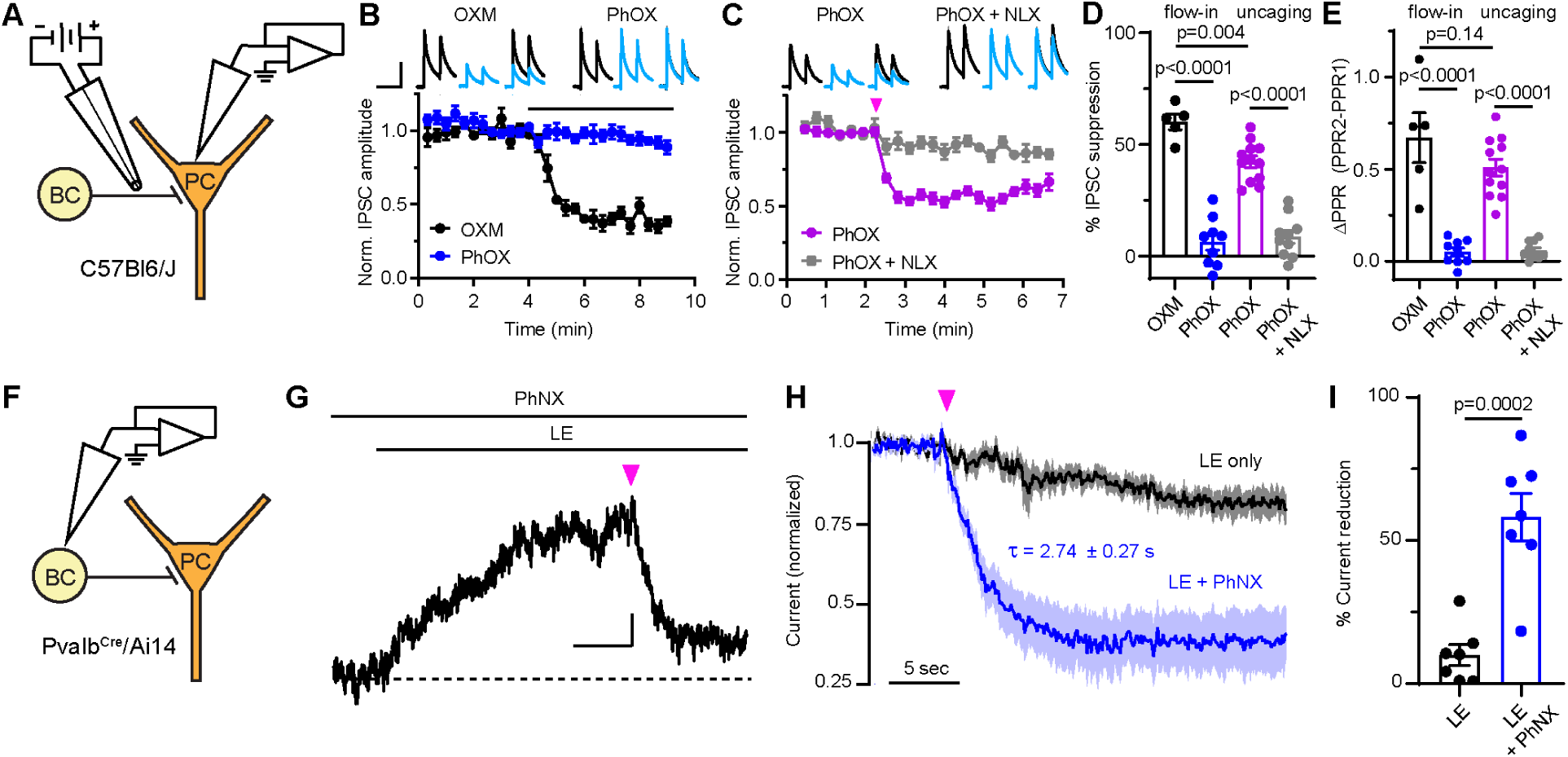
Evaluation of PhOX-mediated photo-agonism and PhNX-mediated photo-antagonism of endogenous opioid receptors in acute hippocampal brain slices. **(A)** Schematic depicting whole-cell voltage clamp recording of opioid-sensitive synaptic inhibition in the CA1 region of hippocampus. A small bipolar stimulating electrode constructed from theta glass was placed near the recorded pyramidal cell (PC) to preferentially stimulate basket cell (BC) axons. **(B)** Baseline-normalized inhibitory post-synaptic currents (IPSCs) in response to bath application of oxymorphone (OXM, =3 µM) or PhOX (3 µM), at the time indicated by the black line (OXM: n=5 cells from 2 mice; PhOX: n=9 cells from 6 mice). Top insets: example average IPSCs (n=3 sweeps from one cell) before (black) and after (blue) application of the indicated drug. Scale bar: 200 pA, 80 ms. **(C)** IPSC suppression in response to photorelease of PhOX (3 µM) with a 50 ms full-field flash of UV light applied at the time indicated by the pink arrow, in the absence or presence of naloxone (NLX, 3 µM) (PhOX: n=12 cells from 7 mice; PhOX + NLX: n=10 cells from 5 mice). Insets are presented as in B. **(D)** Summary data for B and C (One-way ANOVA, F(3,32)=58.2, p<0.0001, Bonferroni’s multiple comparisons test). **(E)** Change in paired pulse ratio (PPR) in response to drug addition and uncaging. Bath application of OXM and PhOX photorelease both produced a significant increase in PPR. (One-way ANOVA, F(3,32)=36.17, p<0.0001, Bonferroni’s multiple comparisons test). **(F)** Schematic depicting the whole-cell voltage clamp recording of opioid-evoked outward currents in CA1 parvalbumin (PV)- basket cells, identified in *Pvalb^Cre^*, Ai14 mice. **(G)** Example photoinhibition of the current evoked by bath application of leucine-enkephalin (LE) upon PhNX uncaging. Scale bar: 10 pA, 10 s. **(H)** Normalized response of LE-evoked currents to a UV light flash in the absence and presence of PhNX (LE only: n=7 cells from 3 mice; LE + PhNX: n=7 cells from 3 mice, plotted as mean ± SEM). **(I)** Summary of the LE-evoked current remaining 10-15 s after application of a light flash in the absence and presence of PhNX (unpaired two-tailed t-test). Current decay in the absence of PhNX is attributed to desensitization. All data are plotted as mean ± SEM.

To validate PhNX, we obtained whole-cell voltage clamp recordings from fluorescently identified parvalbumin-expressing basket cells in hippocampal brain slices from *Pvalb^Cre^*/*Rosa26-lsl-tdTomato* (Ai14) mice. In the presence of PhNX (3 µM), bath application of the mixed mu/delta opioid receptor peptide agonist [Leu^5^]-enkephalin (LE, 300 nM) evoked outward currents that were rapidly reversed by a brief flash of 365 nm light (200 ms, 5 mW) (**Figure 3F-G**). Whereas LE in the absence of PhNX showed only slow desensitization after a light flash, PhNX photoactivation suppressed the LE-evoked current by ∼60% with a time constant of ∼3 sec (**Figure 3H, I**). As with PhOX, PhNX did not affect IPSC amplitude and PPR in the absence of light (**Figure S3**). Collectively, these experiments demonstrate that PhOX and PhNX are inert in the dark and that their photoactivation engages endogenous opioid receptors to modulate neuronal physiology within seconds of light exposure.

### *In vivo* photoactivation after systemic administration of PhOX and PhNX

We next determined whether PhOX and PhNX cross the blood brain barrier (BBB), which is presumably required for effective *in vivo* uncaging. Although the DMNPE caging group lacks functional groups that impart a net charge, in principle it could impede BBB penetrance or produce impractically rapid metabolic clearance. Following retro-orbital intravenous (*i.v.*) administration of equimolar doses of either OXM (5 mg/kg), PhOX (8.5 mg/kg), NLX (10 mg/kg), or PhNX (14 mg/kg) in male and female mice, the concentrations of each drug were determined in the whole brain as well as blood plasma to obtain a brain/plasma ratio as a metric of BBB penetrance. As shown in **Figure 4A**, both PhOX and PhNX exhibited similar brain/plasma ratios to their parent drugs, which suggests that the DMNPE caging group does not impede BBB penetrance and that both PhOX and PhNX should reach useful concentrations in the brain upon systemic administration. We also determined that PhOX (8.5 mg/kg) exhibits a half-life of 90 minutes in the blood after intraperitoneal (*i.p.*) administration (**Figure 4B**), which indicates that *in vivo* uncaging experiments with PhOX can likely be performed for at least an hour after drug administration. Notably, the HCl salts of both PhOX and PhNX are water soluble at millimolar concentrations and could be administered in saline without the addition of surfactants or co-solvents.

**Figure 4.**
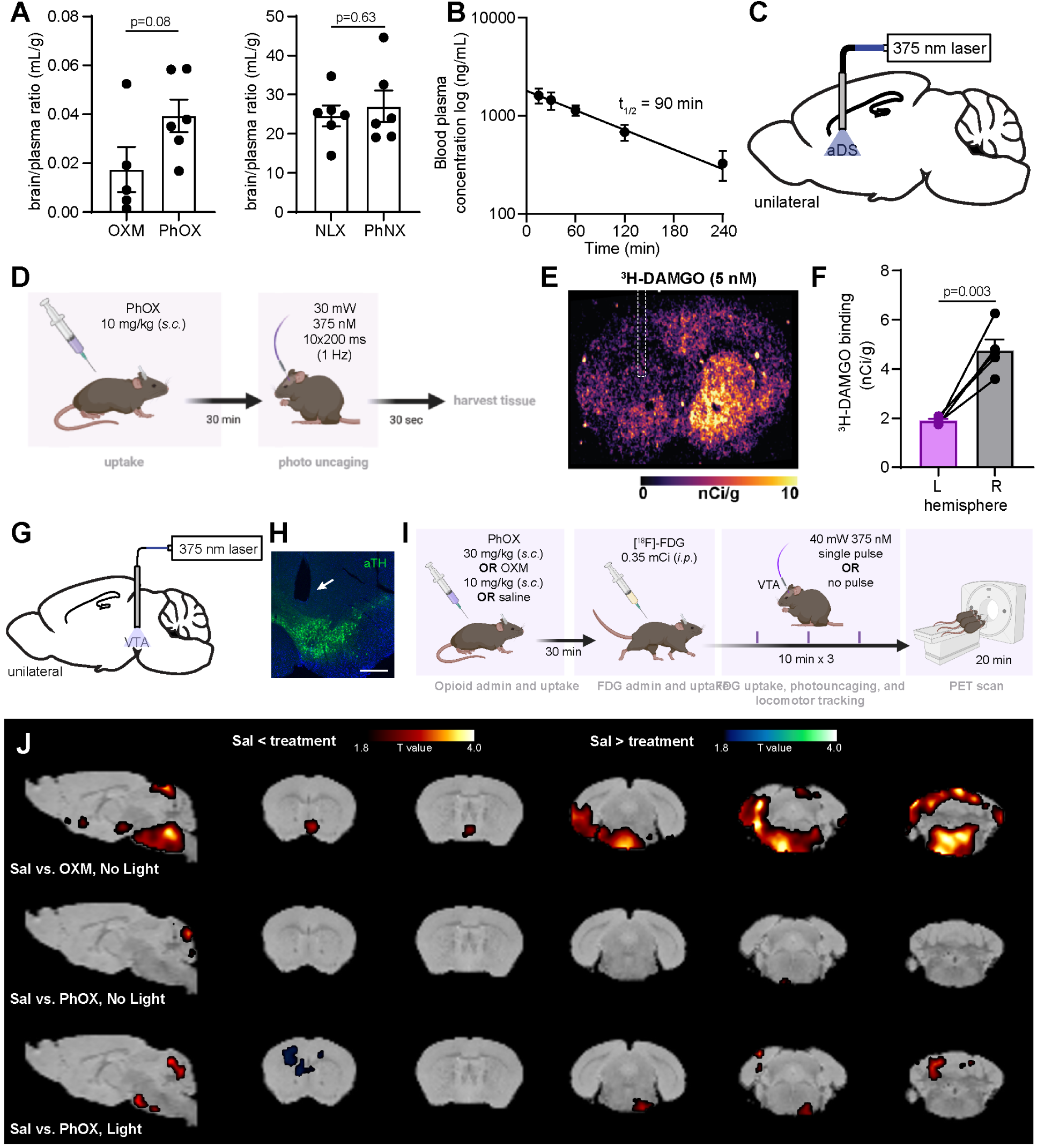
I*n vivo* photoactivation of PhOX after systemic administration. **(A)** Brain-to-plasma ratios of OXM and PhOX (left) and NLX and PhNX (right) determined 15 min after *i.v.* administration (n=5-6 mice per condition, Unpaired two-tailed t-test). **(B)** PhOX presence in the bloodstream after *i.p.* administration (15 mg/kg, n=4 mice). **(C)** Schematic indicating the unilateral implantation of a 375 nm laser-coupled optical fiber in the anterior dorsal striatum. **(D)** Experimental timeline for *in vivo* uncaging followed by autoradiography. **(E)** Autoradiographic image of the fiber implant site in mice treated with PhOX followed by illumination with 375 nm light. **(F**) Quantification of ^3^H-DAMGO binding in the illuminated (L) and unilluminated (R) hemispheres (n=5 sections from 5 mice, paired two-tailed t-test) Data are plotted as mean ± SEM. **(G)** Schematic indicating the unilateral implantation of a 375 nm laser-coupled optical fiber above the ventral tegmental area. **(H)** Representative fluorescence image of the optical fiber implant site in the VTA (scale bar=0.5 mm). **(I)** Schematic depicting the experimental protocol for PhOX uncaging followed by detection of the evoked metabolic activity using small animal PET and [^18^F]-FDG. The mice were anesthetized and scanned 30 min after [^18^F]-FDG was administered, providing a “snapshot” of metabolic activity during the awake, freely-moving state. **(J)** Brain-wide voxel-based analysis was used to evaluate differences in [^18^F]FDG uptake. Color shaded areas overlaid on the corresponding sections of a brain atlas (Ma *et al*., 2005) represent clusters of voxels (≥100) with significant (p < 0.05) increases or decreases in FDG accumulation compared to saline infusions (n=5-6 mice per condition). The contrasts represented are saline vs. OXM (top), saline vs. PhOX without laser stimulation (middle), saline vs. PhOX with laser stimulation (bottom).

To determine if *in vivo* uncaging can be achieved after systemic administration of the caged drug, we turned to *ex vivo* autoradiography, which can reveal receptor occupancy after *in vivo* drug binding to endogenous receptors. To this end, male and female mice were implanted unilaterally with optical fibers in the anterior dorsal striatum (aDS) where MORs are prominent (**Figure 4C**). One week after recovery from surgery, the optical fibers were connected to a 375 nm laser and mice were administered either saline or PhOX (10 mg/kg, *s.c.*), followed 15 min later by application of a series of brief light flashes through the fiber (10 x 200 ms, 1 Hz, 30 mW, **Figure 4D**). Immediately after illumination, mice were anesthetized so that their brains could be removed and flash-frozen within 1-2 minutes of uncaging. Brains were sectioned proximal to the implant site and exposed to a brief (10-minute) incubation with [^3^H]-DAMGO (5 nM) prior to development on a phosphor screen. Quantification of [^3^H]-DAMGO binding revealed that PhOX photoactivation resulted in significant MOR occupancy, as evidenced by decreased [^3^H]-DAMGO binding at the site of illumination compared to the contralateral hemisphere (**Figure 4E,F**). These results demonstrate the feasibility of local *in vivo* drug uncaging using systemic administration and brief flashes of light.

### PET imaging reveals spatially restricted alterations of brain metabolic activity

To assess the spatial control afforded by *in vivo* uncaging, we imaged the induction of metabolic activity in the brain in response to unilateral PhOX photorelease in the ventral tegmental area (VTA, **Figure 4G, H**), where MOR agonists activate mesolimbic dopamine signaling through the disinhibition of dopamine neurons (Gysling and Wang, 1983; Di Chiara and Imperato, 1988a; Johnson and North, 1992). Using small animal positron emission tomography (PET), we compared the brain-wide neural activity evoked by PhOX photorelease in the VTA to systemic OXM administration. As schematized in **Figure 4I**, unilaterally fiber-implanted mice were administered either OXM (10 mg/kg, *s.c.*), PhOX (30 mg/kg, *s.c.*), or saline (*s.c.*), followed 30 min later by [^18^F]-fluorodeoxyglucose (FDG, 0.35 mCi, *i.p.*) to label metabolically active cells. All mice were subsequently connected to a fiber optic cable and placed in a small open field chamber equipped for video monitoring of locomotor behavior. Over the next 30 minutes, half of the saline-treated and PhOX-treated mice were then exposed to 3 x 375 nm light flashes (30 mW, 1 flash every 10 min), whereas the other half received no light. Mice were then anesthetized and placed in the PET scanner to obtain whole-brain maps of metabolic activity during the uncaging period.

Using a significance threshold of p<0.05, the whole brain maps of FDG uptake were compared across saline and drug conditions (**Figure 4J**). Whereas OXM caused widespread increased metabolic activity in opioid-sensitive neural circuits, including the ventral midbrain, and a broad swath of brainstem, PhOX administration in the absence of light minimally altered FDG uptake compared to saline. In contrast, photoactivation of PhOX in the left VTA enhanced FDG uptake in a tightly confined region encompassing the right VTA. This unexpected contralateral signal may reflect compensatory circuit activity related to the uncaging-induced locomotor activation or poorly understood contralateral connectivity in the midbrain. Nonetheless, changes in FDG uptake were restricted to the midbrain in one hemisphere only indicating that the uncaging-induced changes were tightly restricted in space. In all test conditions modest levels of regional cerebellar activation were observed, but the subregion was inconsistent. These results demonstrate that, in the absence of light, PhOX produces minimal changes in brain metabolic activity, whereas PhOX uncaging leads to spatially confined activation of opioid-sensitive neural circuits. These results also raise questions about how local opioid signaling in the VTA may trigger predominantly contralateral metabolic activity.

### *In vivo* PhOX photoactivation drives rapid changes in behavior

To explore the feasibility of using *in vivo* uncaging to study behavior, we asked whether PhOX photoactivation in the brain modulates behavioral responses to pain in mice. We first confirmed that PhOX is inactive in the absence of illumination using a hot plate assay of thermal nociception (**Figure S4A,B**). Compared to saline, PhOX (15 mg/kg, *s.c.*) administration did not alter nocifensive paw withdrawal latencies, nor did it alter the efficacy of a modest dose of morphine (5 mg/kg, *i.p.*).

MOR signaling in the ventral striatum is implicated in pain modulation (Franklin, 1989; Manning, Morgan and Franklin, 1994). We therefore asked whether PhOX photoactivation in the nucleus accumbens medial shell (NAc-mSh) produces analgesia in the Hargreaves thermal pain assay, in which an infrared beam is applied to the hindpaw until withdrawal is observed. (**Figure 5A**). At least one week after unilaterally implanting male and female mice with an optical fiber over the NAc-mSh (**Figure 5B**), we administered PhOX (12 mg/kg *s.c.*), and repeatedly measured withdrawal latencies at the paw contralateral to the implanted fiber. After acquiring a baseline measurement, we applied a thin layer of capsaicin cream (0.1%) to sensitize the hindpaw to the noxious thermal stimulus. Subsequent application of a train of brief 375 light flashes (10 x 200 ms, 1 Hz, 45 mW) transiently returned the paw withdrawal latency to baseline 5 min after light exposure (**Figure 5C**). As a negative control, the same mice were assayed without illumination. Notably, PhOX uncaging in the NAc-mSh did not cause a change in locomotor behavior that might account for the increased latency (**Figure S4C,D**). These results establish that *in vivo* drug uncaging can alter behavior and reveal that opioid signaling in the NAc-mSh can reduce the thermal hypersensitivity produced by capsaicin.

**Figure 5.**
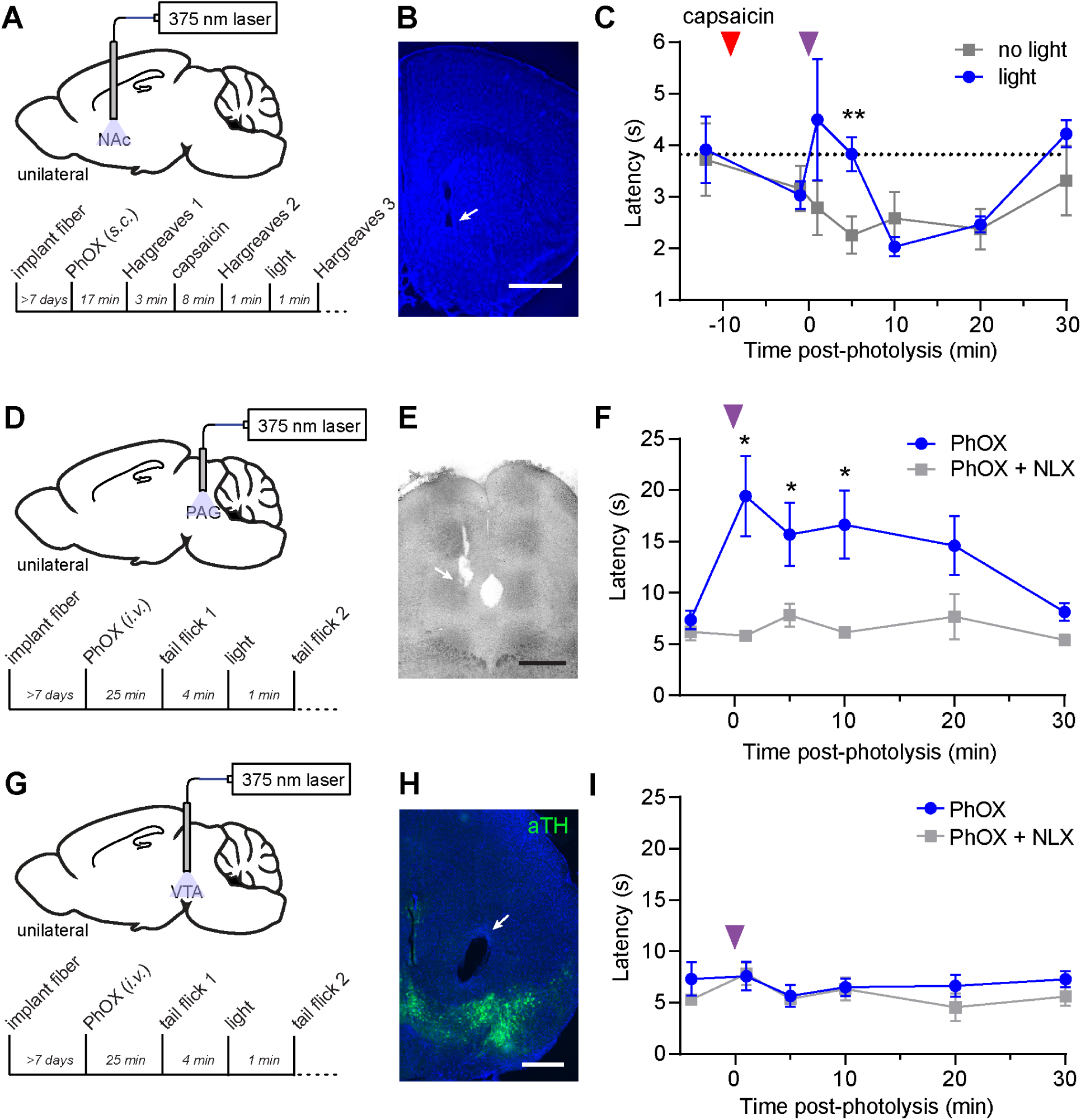
*In vivo* photoactivation of PhOX in the nucleus accumbens and periaqueductal gray suppress pain-related behavior. **(A)** Schematic indicating the unilateral implantation of a 375 nm laser-coupled optical fiber in the nucleus accumbens medial shell (top) and experimental timeline (bottom). **(B)** Representative fluorescence image of the optical fiber implant site (scale bar=1 mm). **(C)** Paw withdrawal latency in the Hargreaves assay in response to PhOX photoactivation in the contralateral NAc-mSh. (n=6 mice, ** indicates p<0.005, paired two-tailed t-test). **(D)** Same as A for the periaqueductal gray. **(E)** Representative brightfield image the optical fiber implant site (scale bar=0.5 mm). **(F)** Withdrawal latency in the tail flick test in response to photostimulation with 375 nm light in the PAG after *i.v.* administration of PhOX or PhOX+NLX (n=6 mice, * indicates p<0.05, Wilcoxon signed-rank test). All data are plotted as mean ± SEM. **(G)** Same as A for the ventral tegmental area. **(E)** Representative fluorescence image of the optical fiber implant site in the VTA (scale bar=0.5 mm). **(I)** Withdrawal latency in the tail flick test before and after unilateral PhOX photoactivation in the VTA. (n=4 mice, no significant differences detected, Mann-Whitney test).

We next attempted PhOX uncaging in the periaqueductal gray (PAG), a brain region in which MOR activation produces strong analgesia in assays of nociception (Yaksh, Yeung and Rudy, 1976) (**Figure 5D**). After recovery from unilateral implantation of an optical fiber in the PAG (**Figure 5E**), male and female mice were administered either PhOX (15 mg/kg, *i.v.*) or PhOX + NLX (PhOX: 15 mg/kg, NLX: 10 mg/kg, *i.v.*) followed by connection of the optical fiber to a 375 nm laser. Beginning 25 minutes after drug administration, nocifensive behavior was monitored using a tail-flick assay. Four minutes after recording a baseline tail-flick latency, a train of brief 375 nm light flashes was applied (10 x 200 ms, 1 Hz, 45 mW). One minute after illumination, tail-flick latency was significantly elevated in mice administered PhOX compared to mice administered both PhOX and NLX (**Figure 5F**). Periodic monitoring revealed that the tail-flick latencies remained elevated for at least 10 minutes after uncaging. In contrast, PhOX photorelease in the VTA, 1.5-2 mm anteroventral to the PAG, did not alter tail-flick latency (**Figure 5G-I**). Together, these results validate the viability of using PhOX uncaging to study pain modulation and indicate that *in vivo* drug uncaging provides good spatial and temporal control over drug action.

To probe the spatiotemporal advantages of drug uncaging in greater detail, we examined behavioral responses to PhOX uncaging in the VTA. MOR signaling in the VTA has multiple behavioral consequences in mice, including increased locomotor activity and behavioral reinforcement (Joyce and Iversen, 1979; Olmstead and Franklin, 1997). To obtain a measure of the *in vivo* uncaging response with temporal resolution on the timescale of seconds, we analyzed the locomotor response to PhOX uncaging in the VTA. After either PhOX (15 mg/kg, *i.v.*) or PhOX + NLX (PhOX: 15 mg/kg, NLX: 10 mg/kg, *i.v.*) administration, mice were connected to the 375 nm laser via a commutator to minimize locomotor impedance, and placed in an open field chamber (**Figure 6A-C**). After 15 minutes of exploration, application of a train of light flashes (10 x 200 ms, 1 Hz, 30 mW) produced a rapid increase in locomotion in mice treated with PhOX but not PhOX + NLX. Strikingly, further analysis of locomotion around the illumination period revealed that locomotion increased within 5 seconds after the first light flash and peaked within 5 seconds after the last flash (**Figure S5**). Similar to the PhOX uncaging-induced analgesia, the resulting locomotor activation persisted for at least 15 minutes after photoactivation. The total distance traveled in the 15 minutes before and after uncaging for each condition is summarized in **Figure 6C**. These results reveal an extremely rapid behavioral response to drug action that supports the use of photopharmacology to drive mechanistic studies *in vivo*.

**Figure 6.**
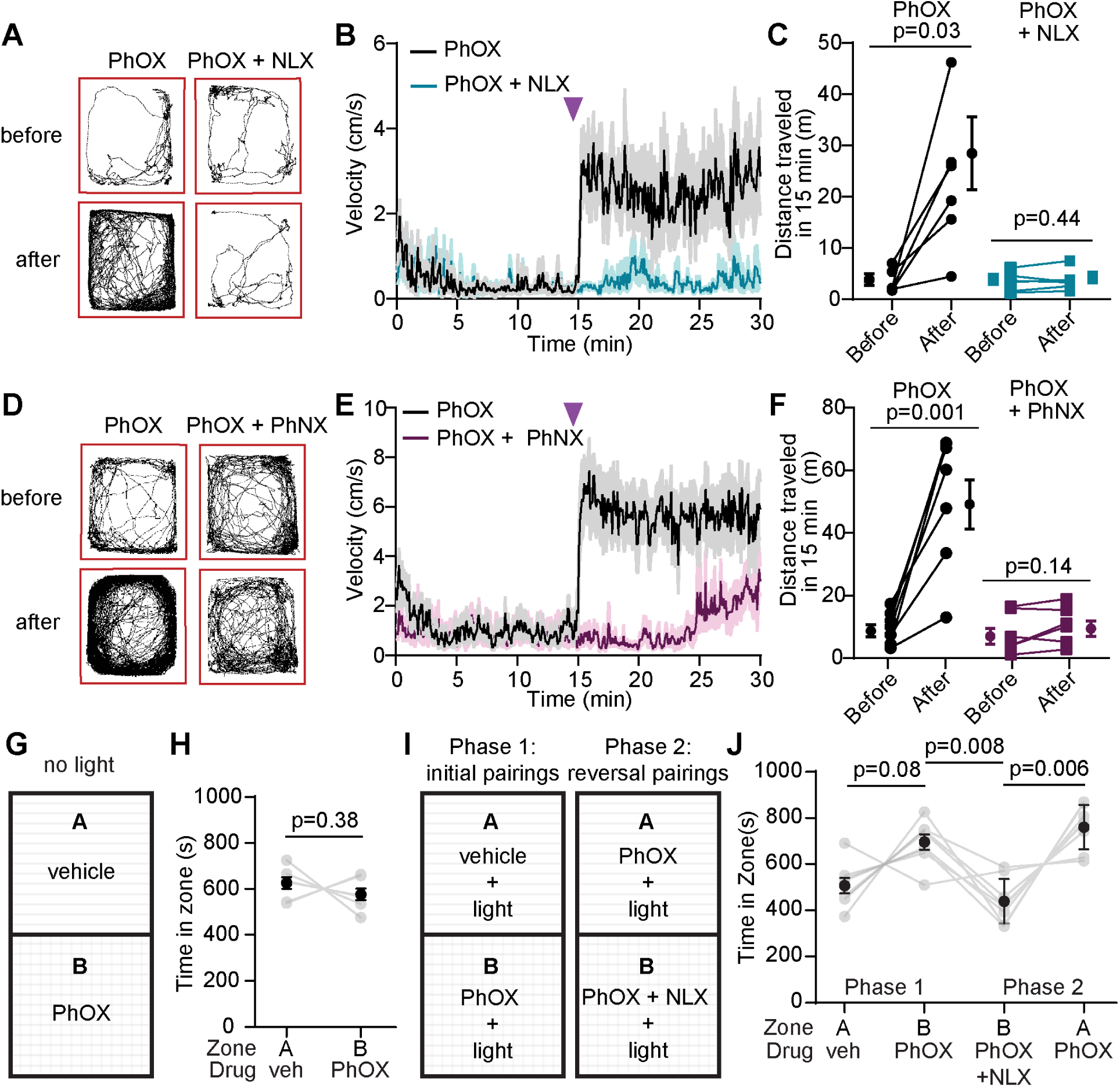
*In vivo* uncaging of PhOX in the ventral tegmental area causes locomotor activation and behavioral reinforcement. **(A)** Example map of open field locomotor activity in response to photostimulation with 375 nm light after *i.v.* administration of PhOX or PhOX+NLX. **(B)** Average plot of velocity over time (n=6 mice). **(C)** Summary plot of the total distance traveled in the 15 minutes before or after photoactivation (n=6 mice, Wilcoxon signed-rank test). **(D)** Example maps of open field locomotor activity in response to photostimulation with 375 nm light after *i.v.* administration of PhOX or PhOX+PhNX. **(E)** Average plot of velocity over time (n=7 mice). **(F)** Summary plot of the total distance traveled in the 15 minutes before or after photoactivation (n=7 mice, paired two-tailed t-test). **(G)** Schematic indicating condition place preference (CPP) context-stimuli pairings without photostimulation. **(H)** Time spent in each zone on test day (n=8 mice, paired two-tailed t-test). **(I)** Schematic indicating CPP context-stimuli pairings and their reversal over two phases of conditioning. **(J)** Time spent in each zone on test day (n=8 mice, Repeated measures one-way ANOVA, F(1.75,12.25)=15.2, p=0.0006, Sidak’s multiple comparison’s test). All data are plotted as mean ± SEM.

To evaluate the potential for *in vivo* uncaging of PhNX, we repeated this experiment in a separate cohort of mice but replaced NLX with PhNX (30 mg/kg), such that the 375 nm light flash would simultaneously release both OXM and NLX and result in light-driven competitive antagonism. As shown in **Figure 6D-F**, inclusion of PhNX completely prevented the initial locomotor response to PhOX photoactivation in the VTA, although 10 min after co-uncaging locomotion began to increase, which suggests that NLX may be cleared more rapidly than OXM after photorelease.

To evaluate the feasibility of using low-cost LED light sources for PhOX uncaging, we compared the locomotor activity elicited using the 375 nm laser, a 365 nm LED, and a 385 nm LED. Although both LEDs were power limited after fiber coupling (∼2 mW), 10 x 200 ms flashes from both LEDs successfully evoked locomotor activity, albeit to a lesser extent than the 375 nm laser set to 15 mW (**Figure S6**). These results demonstrate that modest photoresponses can be obtained using low-cost LEDs, but that a high powered light source such as a UV laser that provides >10 mW of output from the fiber is required to achieve a high degree of PhOX uncaging *in vivo*. Using the locomotor assay, we further explored relationships between the timing and dose of PhOX administration, as well as the number of light flashes (**Figures S7-S8**). These results revealed that photoactivation experiments can begin as early as 15 min after PhOX (s.c.) administration (**Figure S7**), that even the smallest dose tested (2.5 mg/kg, *s.c.*) is sufficient to produce a behavioral response, and that 3-10 flashes (200 ms, 1 Hz, 30 mW) produces the largest response (**Figures S8**).

**Figure 7.**
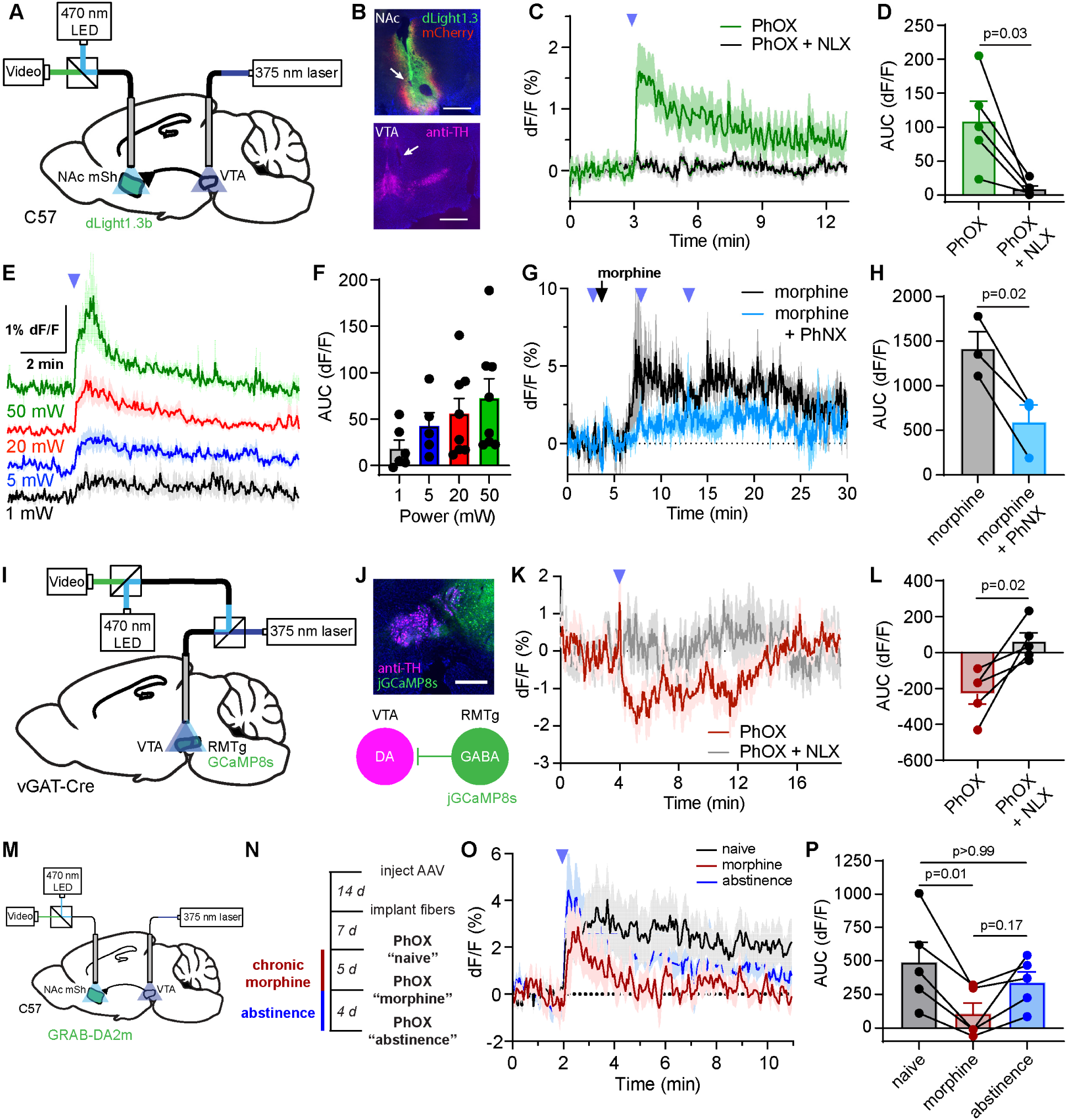
Combined *in vivo* uncaging and fiber photometry reveals drug-dependent changes in the opioid-sensitivity of mesolimbic dopamine signaling. **(A)** Schematic indicating the implantation of a 375 nm laser-coupled fiber in the VTA and a fiber photometry-coupled fiber in the NAc-mSh to detect extracellular dopamine with dLight1.3b in response to PhOX uncaging. **(B)** Example images verifying the sites of photometry fiber implantation along with dLight1.3b (green) and mCherry (red) expression in the NAc-mSh, and uncaging fiber implantation in the VTA by immunostaining for tyrosine hydroxylase (TH, magenta). Scale bars=0.5 mm (NAc) and 0.75 mm (VTA). **(C)** Average dLight1.3b fluorescence (dF/F) in the NAc-mSh in response to PhOX uncaging with a single light flash in the ipsilateral VTA after systemic injection of either PhOX or PhOX + NLX (n=5 mice). **(D)** Summary plot of the uncaging-evoked dlight1.3b fluorescence changes shown in (C) (AUC = area under the curve, n=5 mice, paired two-tailed t-test). **(E)** Average dLight1.3b fluorescence in response to a single light flash at the indicated light power (n=6-8 mice). **(F)** Summary plot of uncaging-evoked dlight1.3b fluorescence changes shown in (E) (n=6-8 mice). **(G)** Average dLight1.3b fluorescence in the NAc-mSh in response to systemic morphine administration with or without subsequent PhNX uncaging in the ipsilateral VTA (n=3 mice). **(H)** Summary plot of the uncaging-evoked dlight1.3b fluorescence changes shown in (G) (n=3 mice, paired two-tailed t-test). **(I)** Schematic indicating the implantation of single fiber in the VTA coupled to both a 375 nm laser and a fiber photometry recording system to detect changes in Ca^2+^ activity with jGCaMP8s expressed in RMTg GABA neurons. **(J)** (Top) Example image of the fiber implant site along with jGCaMP8s expression in RMTg GABA neurons in *Slc32a1*-Cre mice. Scale bar=0.5 mm. (Bottom) Diagram depicting inhibition of VTA dopamine (DA) neurons by jGCaMP8s-expressing RMTg GABA neurons. **(K**) Average normalized jGCaMP8s fluorescence in the VTA in response to PhOX uncaging with a single light flash in the absence and presence of naloxone (n=5 mice). **(L)** Summary plot of the uncaging-evoked jGCaMP8s fluorescence changes shown in (K) (n=5 mice, paired two-tailed t-test). **(M)** Schematic indicating the implantation of a 375 nm laser-coupled fiber in the VTA and a fiber photometry-coupled fiber in the NAc-mSh to detect extracellular dopamine with GRABDA-2m in response to PhOX uncaging. **(N)** Experimental timeline for fiber photometry recordings of VTA PhOX uncaging-evoked dopamine release in the NAc-mSh during chronic morphine administration and abstinence. **(O)** Average GRAB-DA2m fluorescence in the NAc-mSh in response to PhOX uncaging before, during, and after chronic morphine administration (n=5 mice). **(P)** Summary plot of the uncaging-evoked GRAB-DA2m fluorescence changes shown in (C) (n = 5 mice, Friedman test, p=0.0085, Dunn’s multiple comparisons test). All data are plotted as mean ± SEM.

We next asked whether PhOX uncaging in the VTA was able to produce opioid-mediated behavioral reinforcement using a conditioned place preference (CPP) assay. We first established that PhOX is not intrinsically rewarding or aversive in the absence of light by pairing the administration of either PhOX (15 mg/kg, *i.v.*) or vehicle (saline, *i.v.*) in chambers distinguished by visual and tactile cues in a counterbalanced manner (**Figure 6G**). After four conditioning sessions (2 per condition, 20 min in each chamber per session), mice were allowed to roam freely between chambers on the test day. Quantification of the time spent in each zone revealed a lack of preference for either chamber in response to drug administration alone (**Figure 6H**). The effect of light administration was examined in a separate cohort of mice in a two-phase experiment (**Figure 6I**). During Phase 1, after administration of either vehicle or PhOX (15 mg/kg, *i.v.*), mice were connected to the 375 nm laser via a commutator so that light flashes (10 x 200 ms, one bout every 5 min, 20 mW) could be applied during conditioning. On the test day mice were given access to both chambers without being connected to the laser. In all but one mouse, this conditioning paradigm caused the mice to spend a greater amount of time in the PhOX-paired chamber than the vehicle-paired chamber (**Figure 6J**). To confirm that the resulting CPP involves opioid receptor activation, in Phase 2 mice were administered both PhOX and NLX (PhOX: 15 mg/kg, NLX: 10 mg/kg, *i.v.*) in the chamber previously paired with PhOX, and PhOX + vehicle (PhOX: 15 mg/kg, saline, *i.v.*) in the chamber previously paired with vehicle. This pairing reversal led to a strong preference for the PhOX-paired chamber over the PhOX + NLX-paired chamber, indicating that the PhOX-uncaging-driven CPP is due to opioid receptor activation, rather than a side effect of light administration.

### Photopharmacology interfaces with fiber photometry

Photopharmacology offers the opportunity to integrate pharmacological manipulations into experimental frameworks for all-optical interrogation of neural circuits (Packer *et al*., 2015). To explore this possibility while further characterizing the precision of circuit modulation afforded by drug uncaging, we combined PhOX uncaging with fiber photometry. Opioid-evoked locomotor activation and behavioral reinforcement in the VTA are attributed to dopamine release in the nucleus accumbens (NAc) (Leone, Pocock and Wise, 1991; Corre *et al*., 2018). To achieve an optical readout of the physiological response to drug uncaging, we monitored extracellular dopamine in the NAc medial shell (mSh) using the genetically encoded dopamine sensor dLight1.3b while photoactivating PhOX in the VTA (**Figure 7A-B**). After transduction of dLight1.3b along with mCherry as a control for movement artifacts in the left NAc-mSh, mice were implanted with two fibers: one over the left VTA for PhOX uncaging, and another in left NAc-mSh to measure dopamine release. After recovery, mice were administered either PhOX (15 mg/kg, *i.v.*) or PhOX + NLX (PhOX: 15 mg/kg, NLX: 10 mg/kg, *i.v.*), fitted with optical fibers, and then placed into a small open field chamber. After a 10 min baseline period, photoactivation with a single light flash (200 ms, 50 mW) produced a large, rapid increase in extracellular dopamine in the NAc-mSh that was absent in mice co-administered NLX (**Figure 7C-D**). These responses were apparent in the z-score as well (**Figure S9A,B**). The increase in dopamine began within 3 seconds of the light flash, reached 90% of the maximum value within 10 seconds (tau on = 4.25 sec) (**Figure S9C**), and decayed over the course of several minutes (tau off = 5.8 min). Importantly, this optical stimulus failed to produce a dopamine response in the absence of PhOX (**Figure S9D**), indicating that it does not activate dopamine neurons directly (*e.g.* due to tissue heating, (Owen, Liu and Kreitzer, 2019)). Furthermore, application of a strong blue light flash (473 nm, 200 ms, 50 mW) in the presence of PhOX did not evoke detectable dopamine release, verifying that the blue light used for imaging does not lead to significant uncaging (**Figure S9E**). Photoactivation of PhOX (15 mg/kg, *i.v.*) using single flashes of various intensities revealed that 5-20 mW of 375 nm illumination yields reliable, sub-saturating dopamine responses (**Figure 7E-F**). Notably, 1 mW was sufficient to produce a detectable increase in dopamine, indicating relatively high light sensitivity under these PhOX administration conditions. Application of two light flashes separated by several minutes drove dopamine release repeatedly (**Figure S9F**). These results demonstrate the compatibility of *in vivo* photopharmacology with fluorescence-based measurements of neural function and establish a robust, spatiotemporally precise assay for probing the opioid-sensitivity of mesolimbic dopamine circuitry.

Using the same experimental configuration, we assessed the ability of unilateral PhNX uncaging to suppress the activation of the mesolimbic dopamine circuit by systemic morphine (**Figure 7G-H**). As expected, morphine (10 mg/kg, *i.p.*) injection produced a large increase in NAc-mSh dopamine release that lasted for at least 30 minutes. However, photoactivation of PhNX in the ipsilateral VTA (15 mg/kg, *i.v.*, injected 15 min prior to morphine, 20 x 200 ms, 1 Hz, 80 mW) just before and after morphine injection attenuated this dopamine increase. This result further validates the use of PhNX for *in vivo* applications.

Thus far, we photoactivated opioid drugs and obtained fluorescence measurement of the physiological response at discrete sites, which minimizes experimental confounds due to sensor bleaching and/or photoconversion between photostates that may result from illuminating fluorescence probes with ultraviolet light. Yet photoactivation at the recording site enables the unequivocal assignment of experimental outcomes to receptor engagement in that brain region. For example, MOR agonists are thought to activate VTA dopamine neurons through a disinhibitory mechanism involving the suppression of GABA release from spontaneously active GABA neurons. *In vivo* and *ex vivo* electrophysiology experiments implicate opioid-sensitive GABA neurons located in the adjacent rostromedial tegmental nucleus (RMTg) (Jalabert *et al*., 2011; Matsui and Williams, 2011; Matsui *et al*., 2014), but this has been difficult to directly observe *in vivo* due to challenges associated with site-specific drug delivery during cell type-specific recording of neural activity. We reasoned that direct measurement of the impact of PhOX photoactivation on Ca^2+^ activity in RMTg GABA neuron terminals in the VTA would be a convenient and advantageous approach to addressing this issue.

To enable photouncaging at the site of fiber photometry recording, we constructed a relay system that feeds the 375 nm laser into the photometry output pathway (**Figure 7I**, **Figure S10**). To directly ask if the output of RMTg GABA neurons is suppressed by opioids in the VTA *in vivo*, we transduced RMTg in male and female *Slc32a1*-Cre (vGAT-IRES-Cre) mice with the genetically-encoded Ca^2+^ indicator jGCaMP8s (Zhang *et al*., 2021) along with mCherry for motion correction and then implanted a single optical fiber over the VTA (**Figure 7J**). This allowed us to record Ca^2+^ activity in RMTg GABA neuron terminals in the VTA while locally photoreleasing OXM from PhOX at the same location. To ask if MOR activation modulates the output from RMTg GABA neurons in the VTA *in vivo*, we applied a single 375 nm light flash (200 ms, 20 mW) while recording green and red fluorescence through the fiber, before and after the administration of PhOX (30 mg/kg, *s.c.*). In the absence of PhOX, the 375 nm light flash caused a transient increase in green fluorescence that decayed over several minutes (tau off = 3.1 min, **Figure S10**). This artifact was consistent within individual mice, which allowed us to correct for it by subtracting the average of 3 pre-injection trials from post-drug administration recordings. This analysis revealed that PhOX (15 mg/kg *i.v.*) photoactivation with a single light flash (200 ms, 20 mW) suppressed intracellular Ca^2+^ signaling in RMTg GABA neuron terminals located within the VTA (**Figure 7K-L)**. Co-administration of NLX (10 mg/kg) blocked the observed suppression, confirming that it is due to opioid receptor activation, as opposed to ineffective correction of the UV light-induced artifact. Together, these results demonstrate the compatibility of *in vivo* uncaging with fiber photometry and reveal that RMTg GABA neuron terminals are suppressed locally by MOR agonists in the VTA *in vivo*.

### *In vivo* opioid uncaging reveals mesolimbic dopamine circuit adaptations to sustained morphine exposure

A primary issue with opioid drugs is their propensity to produce tolerance upon sustained consumption. This greatly complicates their clinical use for pain treatment and contributes to the escalation of drug-seeking behavior in the context of opioid addiction. Although MORs are known to desensitize on multiple timescales in response to sustained activation (Williams *et al*., 2013), how the opioid-sensitivity of the mesolimbic dopamine circuit is altered by prolonged MOR activation is not known. Because PhOX uncaging restricts drug action to the VTA while producing a highly stereotyped, readily observed dopamine transient in the NAc, *in vivo* opioid photopharmacology provides a unique opportunity to probe for changes within the VTA without confounds due to opioid drug action other brain areas, including the NAc (Pentney and Gratton, 1991; Borg and Taylor, 1997; Britt and McGehee, 2008).

After expression of the dopamine sensor GRAB-DA2M and fiber implantation (**Figure 7M**), the NAc dopamine response to VTA MOR activation with PhOX uncaging was measured before, during, and several days after administrating mice escalating doses of morphine (twice a day with 10, 20, 30, 40, and 50 mg/kg over 5 days, **Figure 7N**). Importantly, photometry measurements were taken ∼8 hrs after morphine administration such that any changes observed would not be attributable to occlusion, as the half-lives of morphine and its glucuronide adducts are on the order of 30 min (Handal *et al*., 2002). Compared the morphine-naïve state, chronic morphine treatment attenuated the NAc dopamine response to PhOX uncaging with 375 nm light (10 x 200 ms, 1 Hz, 20 mW). Although the peak responses were similar, the dopamine transient returned to baseline within 3 min of illumination after chronic morphine treatment, whereas it lasted >9 minutes prior to chronic morphine administration (naïve, **Figure 7O**). After several days of abstinence, the NAc dopamine response to VTA MOR activation showed a trend toward recovery. In contrast, chronic saline treatment did not alter the VTA opioid-NAc dopamine coupling in a separate cohort of mice (**Figure S11**). These results indicate that the opioid-sensitivity of the mesolimbic dopamine circuit is reduced during prolonged MOR activation by morphine. Because dopamine signaling in the NAc is strongly implicated in drug reward, this opioid-dopamine decoupling is likely a key factor contributing to the tolerance that underlies increased drug seeking and dose escalation during opioid addiction.

## Discussion

In this study we have demonstrated that *in vivo* photopharmacology interfaces well with temporally resolved behavioral and neurophysiological studies that are commonly used to investigate the mammalian nervous system. Importantly, we showed that careful molecular design can produce water soluble, BBB-penetrant caged drugs that can be effectively released in the brain through implanted optical fibers upon illumination with modest light intensities (200 ms flashes, 1-20 mW) that are well tolerated in optogenetics experiments (Owen, Liu and Kreitzer, 2019). Because the caging group used does not absorb blue and green light, photoactivation could be selectively achieved with UV light during one-photon optical recordings of intracellular Ca^2+^ and extracellular dopamine using green and red fluorophores.

Furthermore, we showed that optical recording and drug photoactivation can be conducted at the same site through the same optical fiber, which provides important experimental advantages. Finally, we showed that the robust stimulus-response relationship afforded by drug photoactivation can be used as the basis of longitudinal studies that track experimentally-induced changes in stimulus-response coupling. In the process of establishing an experimental framework for using photopharmacology to study the brain, we also validated two new reagents (PhOX and PhNX) that can be used to bidirectionally control endogenous MOR signaling *in vivo*.

In response to PhOX photoactivation in the VTA, we reliably observed dopamine release in the NAc using fiber photometry, consistent with numerous *in vivo* studies in rodents (Di Chiara and Imperato, 1988a, 1988b; Spanagel, Herz and Shippenberg, 1990). Yet human studies based on PET imaging of [^11^C]-raclopride displacement from D2 dopamine receptors are more conflicted. Although two studies found no increase in NAc dopamine in response to systemic administration of MOR agonists in heroin addicts (Daglish *et al*., 2008; Watson *et al*., 2014), a more recent study observed morphine-evoked NAc dopamine release in opioid-naïve subjects (Spagnolo *et al*., 2019). Our finding that VTA opioid-evoked NAc dopamine release is diminished with chronic opioid exposure provides a mechanistic explanation for this apparent conundrum. We also found that MOR agonist locally suppresses output from GABAergic RMTg neurons in the VTA using same-site fiber photometry and PhOX photoactivation. In conjunction with *ex vivo* brain slice electrophysiology studies (Matsui *et al*., 2014), our results indicate that diminished opioid-sensitivity of RMTg GABA neuron input to VTA dopamine neurons underlies tolerance in the mesolimbic dopamine pathway. As the VTA is a central site for opioid drug reward, these circuit adaptations are likely to account for the tolerance to the rewarding effects of opioids that occurs after sustained drug use, a hallmark of opioid addiction (Christie, 2008).

Photopharmacology offers numerous advantages over traditional pharmacological methods employed in behavioral pharmacology and optical physiology experiments. Using light, local drug administration can be achieved in a relatively non-invasive manner during ongoing behavior experiments with minimal disruption or artifact generation. Rapid, time-locked drug release provides a robust stimulus to which a response can be readily matched and quantified (*e.g.* amplitude, integral, kinetics). The ability to make observations immediately after drug delivery also minimizes confounds due to receptor desensitization, which occurs on the timescale of milliseconds-to-minutes, depending on the receptor. Conveniently, drug dosage can be easily tuned by varying the amount of light delivered. Finally, photopharmacology is compatible with optical recording methods, and in principle, electrophysiological recording methods (*e.g.* via optrodes). By restricting drug action to the recording site, the observed response can be unequivocally attributed to local drug action as opposed to upstream circuits that are engaged by systemic drug administration.

A primary advantage of using photopharmacology is the ability to deliver a brief pharmacological stimulus that is time-locked to external events under control of the experimenter. Using PhOX, we observed physiological and behavioral responses to photoactivation within seconds of illumination. Using weak stimuli (*e.g.* single light flashes), responses lasted for lasted ∼10 min, despite being small in amplitude (**Figure 7E**). These deactivation kinetics likely result from relatively slow diffusion and metabolic clearance of OXM from the site of photorelease. In some experimental contexts, a time window of several minutes after light application may be ideal (*e.g.* behavioral testing before and after delivery), but it imposes constraints on others (*e.g.* repeated administration). While it may be possible to tune the deactivation kinetics to some extent using different agonists, photoswitchable ligands that can be rapidly inactivated by light (Gorostiza and Isacoff, 2008; Broichhagen, Frank and Trauner, 2015) or in the dark (Mourot *et al*., 2011) are likely to offer a narrower time window of receptor activation.

Our experimental framework for *in vivo* photopharmacology sets the stage for development and validation of other light-sensitive ligands and should extend to other targeted drug-delivery approaches such as well (*e.g.* ultrasound drug uncaging) (Wang *et al*., 2018). This involves establishing BBB penetrance with systemic drug administration, using a non-behavioral readout of *in vivo* photoactivation-mediated target engagement (*e.g.* receptor occupancy, PET), achieving a rapid (*i.e.* seconds-timescale) behavioral response to brief illumination, and concomitant recordings of the neural response to drug action. In particular, we make a case for continued investment in UV-sensitive caging groups that do not perturb BBB penetrance, as they are readily compatible with widespread 1-photon fluorescence methods such as fiber photometry. Because DMNPE caging group and its relatives have poor 2-photon cross sections, they are likely to be compatible with 2-photon imaging as well.

Another key advantage of caged drugs and photoswitchable ligands is that they act on endogenous receptors with anatomically-defined circuit-specificity. However, they do not currently offer cellular specificity. Genetically-targeted tools can enable cell type-specific manipulation of endogenous receptor signaling *in vivo* (DARTs) (Shields *et al*., 2017). As variations based on photoswitches (Lin *et al*., 2015; Donthamsetti *et al*., 2019, 2021) and caged ligands (Tobias *et al*., 2021) continue to evolve, the performance benchmarks established in this study will help guide the development of optimal photopharmacological tools for *in vivo* applications.

The therapeutic potential of *in vivo* photopharmacology should not be overlooked, as light provides a means to achieve spatially targeted drug delivery that minimizes side effects due to drug action at other sites, as well as means to restrict drug action only to the time it is needed. These features are particularly desirable in the context of opioid analgesics, which, in addition to suppressing pain at multiple sites throughout the brain and spinal cord, can have undesirable adverse effects such as severe respiratory depression and constipation, and are extremely addictive. Systemically available photoactivatable opioids such as PhOX open the door to the tantalizing possibility of restricting opioid action to sites that produce pain relief, only when pain is felt or anticipated, using an external or implanted light source.

## Acknowledgements

We thank the National Institute on Drug Abuse Drug Supply Program (NDSP) for generously providing pharmacological reagents; A. X. and J. Momper for contributing to bioavailability experiments; G. Or and L. Tian for dLight1.3b AAV; Y. Li for GRABDA-2m; E. Berg for genotyping, animal husbandry, adenoassociated virus production and administrative assistance; and T. Gremel, K. Tye, T. Hnasko, and B.K. Lim for helpful discussions. The project was supported by the Brain & Behavior Research Foundation (M.R.B.), the Esther A. & Joseph Klingenstein Fund & Simons Foundation (M.R.B.), NIH grants R00DA034648 (M.R.B), U01NS113295 (M.R.B. & Diversity Supplements to D.A.J. and A.E.L.), T32GM007240 (X.J.H.), ZIA000069 (M.M.), and Plan Nacional Sobre Drogas, Ministerio de Sanidad, Spain 2021I070 (J.B.). Some figures were generated in part using BioRender.

## Author contributions

M.R.B. conceived and coordinated the project with help from S.P.M. and X.M. X.M. synthesized and characterized PhOX, PhNX, and DEAC-OXM with help from J. Z., and performed the *in vitro* characterization of G protein signaling with help from A.E.L. A.R. performed the *in vitro* characterization of β-arrestin signaling with oversight from J.B.. S.P.M. performed and oversaw all aspects of the evaluation of PhOX and PhNX in behaving mice with help from D.A.J., S.T.L., & G.L.. S.P.M., D.A.J. performed analgesia, locomotion, conditioned place-preference, and fiber-photometry experiments. J.C.Y. and E.V. performed histology. S.T.L. and G.L. contributed to analgesia experiments. C.A.J. evaluated PhOX and PhNX using brain slice electrophysiology with help from X.J.H. E.V. performed the autoradiography experiments with help from M.L., J.L.G. & J.B. with oversight from M.M. E.V. and S.P.M. performed the PET imaging experiments with help from M.L. and J.L.G., and J.B. analyzed the data, with oversight from M.M. Each author analyzed the data they generated with oversight from S.P.M., J.B., M.M., and M.R.B. S.P.M., X.M. and M.R.B. wrote the manuscript with input from all authors.

## Ethics statement

Animal experimentation: All procedures were performed in accordance with protocols approved by the University of California San Diego and the National Institute on Drug Abuse Intramural Research Program’s Institutional Animal Care and Use Committees (IACUC) following guidelines described in the US National Institutes of Health Guide for Care and Use of Laboratory Animals (UCSD IACUC protocol S16171, NIDA IACUC protocol 21-NRB-43). All surgery was performed under isoflurane anesthesia.

## STAR Methods

### Resource Availability Lead Contact

Further information and requests for resources and reagents should be directed to and will be fulfilled by the lead contact, Dr. Matthew Banghart.

### Materials availability

Samples of PhOX and PhNX are available upon request.

### Data and code availability

Source data for all figures are available in the supporting information. Raw data are available upon request.

## Experimental model and subject details

### Cell lines

The commercially available HEK293T cell line (sex: female) was used in this study. Detailed growth conditions are reported in the methods details section.

### *Ex vivo* electrophysiology from mouse brain slice

For *ex vivo* brain slice electrophysiology, both male and female wild-type C57/Bl6 or PValb-Cre/Ai14 mice used were postnatal day 15-35 at the time of brain extraction.

### Mice for *in vivo* photoactivation experiments

For *in vivo* photoactivation experiments, 2-6 month old wild-type C57/Bl6 or vGAT-Cre mice were used either from Jackson Labs or bred in house on a C57/bl6j background. Mice were maintained on a reverse light dark cycle (12:12 dark:light) with *ad libitum* access to food and water and nesting material for environmental enrichment. Both male and female mice were used. All experiments were performed during the dark cycle. Animal handling protocols were approved by the UC San Diego Institutional Animal Care and Use Committee.

## Method details

### Chemical synthesis and characterization

Commercial reagents were obtained from reputable suppliers and used as received. Oxymorphone free base was obtained from Noramco (516340913), naloxone HCl was obtained from Sigma Aldrich (N7758), 4-(bromomethyl)-7-(diethylamino) coumarin was obtained from TCI (B5008), and 1-(1-bromoethyl)-4,5-dimethoxy-2-nitrobenzene was synthesized according to a published procedure (Ren, Ji and Ai, 2015). All solvents were purchased as septum-sealed bottles stored under an inert atmosphere. All reactions were sealed with septa through which a nitrogen atmosphere was introduced unless otherwise noted. Reactions were conducted in round-bottomed flasks or septum-capped amber screw-cap vials containing Teflon-coated magnetic stir bars. Room lights were covered with Roscolux Canary Yellow #312 film (Rosco Laboratories, Stamford, CT) to filter out wavelengths of light that could lead to unintentional photolysis during purification and handling.

1-methyl-4,5-dimethoxy-2-nitrobenzene oxymorphone (**PhOX**)

To a stirred solution of oxymorphone (80 mg, 0.265 mmol) and 1-(1-bromoethyl)-4,5-dimethoxy-2-nitrobenzene (92 mg, 0.317 mmol) in anhydrous DMF (1 mL) in an amber glass vial, anhydrous K_2_CO_3_ (73 mg, 0.528 mmol) was added. The mixture was stirred at 22 °C . After 16 h, TLC showed the completion of the reaction and water (1 mL) was added. The mixture washed with EtOAc (5 mL x 2). The combined organic phases were washed with 5% LiCl (10 mL) and brine (10 mL). The organic phase was dried over Na_2_SO_4_ and concentrated under vacuum. The residual was purified by column chromatography (SiO_2_, EtOAc → EtOAc/MeOH (9:1)) to give PhOX as a light-yellow oil (85 mg, 63%).

^1^H NMR (300 MHz, CDCl_3_) *δ* 7.58 (m, 1H), 7.38 – 7.24 (m, 1H), 6.62 – 6.19 (m, 3H), 5.02 (s, 1H), 4.62 (m, J = 11.2 Hz, 1H), 4.05 – 3.87 (m, 6H), 3.12 – 2.88 (m, 2H), 2.81 (m, 1H), 2.52 – 2.30 (m, 6H), 2.26 – 2.15 (m, 1H), 2.06 (m 1H), 1.88 – 1.77 (m, 1H), 1.72 (m, 3H), 1.59 – 1.34 (m, 2H).

^13^C NMR (75 MHz, CDCl_3_) *δ* 208.10, 207.61, 154.25, 153.77, 147.75, 147.61, 145.73, 144.99, 140.32, 140.28, 139.84, 139.76, 135.24, 134.75, 129.78, 129.50, 126.13, 125.17, 119.62, 119.41, 118.47, 115.58, 108.86, 108.53, 107.67, 107.42, 90.41, 90.35, 73.92, 71.81, 70.25, 70.23, 64.48, 64.41, 56.69, 56.42, 56.26, 50.36, 50.10, 45.17, 45.12, 42.66, 36.05, 35.96, 31.50, 31.37, 30.57, 30.37, 23.57, 23.46, 21.93, 21.86.

LR-MS (ESI) *m/z* 511 [(M+H)^+^, 70%], 533 [(M+Na)^+^, 100%].

HR-MS *m/z* 533.1889 [M+Na]^+^ (calcd for C_27_H_30_N_2_O_8_Na, 533.1884).

1-methyl-4,5-dimethoxy-2-nitrobenzene naloxone (**PhNX**)

To a stirred solution of naloxone hydrochloride dihydrate (106 mg, 0.265 mmol) and 1-(1-bromoethyl)-4,5-dimethoxy-2-nitrobenzene (92 mg, 0.317 mmol) in anhydrous DMF (1 mL) in an amber glass vial 22 °C, anhydrous K_2_CO_3_ (110 mg, 0.796 mmol) was added. After 16 h, TLC showed the completion of the reaction and water (1 mL) was added. The mixture was washed with EtOAc (5 mL x 2). The combined organic phases were washed with 5% LiCl (10 mL) and brine (10 mL). The organic phase was dried over Na_2_SO_4_ and concentrated under vacuum. The residual was purified by column chromatography (SiO_2_, hexane → hexane/EtOAc (9:1)) to give **PhNX** as a light-yellow oil (95 mg, 67%).

^1^H NMR (300 MHz, Methanol-d_4_) *δ* 7.58 (m, 1H), 7.34 (m, 1H), 6.78 – 6.50 (m, 2H), 6.35 – 6.23 (m, 1H), 5.88 (m, 1H), 5.20 (m, 2H), 4.71 (m, 1H), 3.91 (m, 6H), 3.26 – 2.84 (m, 5H), 2.65 – 2.30 (m,3H), 2.23 – 1.94 (m, 2H), 1.88 – 1.64 (m, 4H), 1.59 – 1.47 (m, 1H), 1.45 – 1.15 (m, 1H).

^13^C NMR (75 MHz, Methanol-d_4_) *δ* 208.66, 153.81, 147.90, 145.55, 140.15, 139.97, 135.31, 134.40,

129.95, 126.86, 119.48, 119.12, 116.88, 108.82, 107.59, 90.16, 73.92, 70.18, 61.92, 57.19, 55.42, 55.35, 50.62, 43.02, 35.29, 31.29, 30.22, 22.30, 22.22.

LR-MS (ESI) *m/z* 537 [(M+H)^+^, 100%], 559 [(M+Na)^+^, 35%].

HR-MS *m/z* 537.2230 [M+H]^+^ (calcd for C_29_H_33_N_2_O_8_, 537.2231).

diethylaminocoumarin-oxymorphone (**DEAC-OXM**)

To a solution of oxymorphone (7.2 mg, 0.024 mmol) in dry DMF (2 ml) was carefully added compound 4-(bromomethyl)-7-(diethylamino) coumarin (9.2 mg, 0.026 mmol) and K_2_CO_3_ (9.9 mg, 0.072 mmol) at 22 °C. After 16 h, TLC showed the completion of the reaction and water (1 mL) was added. The aqueous layer was extracted with ethyl ether (2 x 10 mL). The combined organic phases were washed with 5% LiCl (10 mL) and brine (10 mL). The organic phase was dried over Na_2_SO_4_ and concentrated under vacuum. The organic phase was dried over Na_2_SO_4_ and concentrated under vacuum. The residual was purified by column chromatography (SiO_2_, hexane → hexane/EtOAc (9:1)) to give DEAC-OXM (10 mg, 81%) as a yellow oil.

^1^H NMR (600 MHz, Methanol-d_4_) δ 7.62 (d, *J* = 9.0 Hz, 1H), 6.83 (d, *J* = 8.2 Hz, 1H), 6.76 (dd, *J* = 9.1, 2.6 Hz, 1H), 6.70 (dd, *J* = 8.2, 0.9 Hz, 1H), 6.55 (d, *J* = 2.6 Hz, 1H), 6.24 (d, *J* = 1.2 Hz, 1H), 5.50 (dd, *J* = 15.0, 1.3 Hz, 1H), 5.38 (dd, *J* = 15.0, 1.3 Hz, 1H), 4.83 (s, 1H), 3.49 (q, *J* = 7.1 Hz, 4H), 3.06 (td, *J* = 14.4, 5.1 Hz, 1H), 2.96 (d, *J* = 5.8 Hz, 1H), 2.62 (dd, *J* = 18.8, 5.8 Hz, 1H), 2.58 – 2.47 (m, 2H), 2.45 (s, 3H), 2.25 – 2.12 (m, 2H), 1.94 – 1.85 (m, 1H), 1.65 – 1.56 (m, 1H), 1.54 – 1.46 (m, 1H), 1.23 (t, *J* = 7.1 Hz, 6H).

^13^C NMR (150 MHz, Methanol-d_4_) δ 209.27, 163.31, 156.13, 152.99, 150.98, 145.32, 140.76, 130.21, 127.34, 125.19, 119.72, 118.80, 109.02, 106.09, 105.02, 96.81, 90.66, 70.36, 67.94, 64.41, 50.16, 45.05, 44.21, 41.53, 35.40, 31.43, 29.90, 21.63, 11.36.

LR-MS (ESI) *m/z* 531 [(M+H)^+^, 100%].

HRMS (m/z): calculated for [C_31_H_35_N_2_O_6_]^+^: 531.2490, found 531.2485.

### RE-DEAC-OXM

At room temperature, 500 mL DEAC-OXM HCl (50 uM in PBS) was stirred vigorously and exposed 405 nm (**13.5 mW**). After 16 h, then the pH of the solution was adjusted to ∼8 and extracted with EtOAc (500 mL x 2). Combined organic layers were washed with brine (500 mL), dried by Na_2_SO_4,_ and concentrated under vacuum. The residue was purified by C18 flash chromatography (water/CH_3_CN, 95%/5%→ 100%) to give **RE-DEAC-OXM** (4 mg, 31%) as a yellow solid.

^1^H NMR (600 MHz, Methanol-d_4_) δ 8.49 (s,1H), 7.63 (d, *J* = 9.1 Hz, 1H), 6.69 (dd, *J* = 9.1, 2.6 Hz, 1H), 6.66 (s, 1H), 6.49 (d, *J* = 2.6 Hz, 1H), 5.71 (s, 1H), 4.10 (d, *J* = 15.9 Hz, 1H), 3.93 (d, *J* = 15.9 Hz, 1H), 3.57 (d, *J* = 5.9 Hz, 1H), 3.45 (q, *J* = 7.1 Hz, 4H), 3.35 (d, *J* = 19.3 Hz, 1H), 3.11 (dd, *J* = 12.7, 4.2 Hz, 1H), 3.00(dd, *J* = 19.3, 5.9 Hz, 1H), 2.84 (s, 3H), 2.72 (m, 2H), 2.26 (dt, *J* = 19.3, 3.2 Hz, 1H), 1.99 (dd, *J* = 14.2, 3.2 Hz, 1H), 1.67 (m, 2H), 1.18 (t, J = 7.1 Hz, 6H).

^13^C NMR (150 MHz, Methanol-d_4_) 210.22, 165.07, 158.82, 157.42, 152.34, 145.65, 139.48, 129.33, 127.60, 127.23, 122.59, 122.13, 110.38, 109.39, 108.09, 98.08, 91.02, 71.53, 68.01, 50.15, 48.35, 45.58, 41.82, 32.32, 28.93, 24.30, 12.75.

LR-MS (ESI) *m/z* 531 [(M+H)^+^, 100%].

HRMS (m/z): calculated for [C_31_H_35_N_2_O_6_]^+^: 531.2490, found 531.2493.

### Authentication and data analysis

Reactions were monitored by liquid chromatography-mass spectrometry (LC-MS) using C-18 column (4.6 × 50 mm, 1.8 μm, Agilent) with a linear gradient (water/MeCN 5%/95% → MeCN 100%, 0-8 min with 0.1% formic acid, 1 ml/min flow, electrospray ionization, positive ion mode, UV detection at 210 nm, 254 nm, and 350 nm). High-resolution mass spectrometry data were obtained at the UCSD Chemistry and Biochemistry Mass Spectrometry Facility on an Agilent 6230 time-of flight mass spectrometer (TOFMS). Proton (^1^H) and carbon (^13^C) NMR spectra were recorded at room temperature in base-filtered CDCl3 on a Bruker AVA-300 spectrometer operating at 300 MHz for proton and 75 MHz for carbon nuclei. For ^1^H NMR spectra, signals arising from the residual protioforms of the solvent were used as the internal standards. ^1^H NMR data are reported as follows: chemical shift (δ) [multiplicity, coupling constant(s) J (Hz), relative integral] where multiplicity is defined as: s = singlet; d = doublet; t = triplet; q = quartet; m = multiplet or combinations of the above. All NMR spectra were processed using MestReNova 14.2.1. UV-visible spectra were recorded on a NanoDrop 2000 UV-VIS spectrophotometer (Thermo-Fisher).

### *In vitro* uncaging and dark stability

To determine dark stability, PhOX and PhNX (1 mM) were dissolved phosphate-buffered saline (PBS, pH 7.2) and left in the dark for 24 h. Comparison of samples taken at 0 and 24 h by HPLC-MS (1260 Affinity II, Agilent Technologies, Santa Clara, CA, USA) revealed no obvious decomposition or conversion to oxymorphone or naloxone. In addition to determining the chemical composition of the uncaging product by HPLC-MS, the initial photolysis rate of PhNX and PhOX was compared using HPLC in response to illumination with 375 nm light. Solutions of PhOX and PhNX (1 mM) dissolved in PBS buffer (pH 7.2) were placed in 1 mL glass vials with stir bars and illuminated at a light intensity of 20 mW from the output of a 375 nm laser (LBX-375-400-HPE-PPA, Oxxius, France) via an optical fiber (FT200UMT, 200 µm, 0.39 NA). The solutions were illuminated in 15 sec periods.

### Data analysis

Samples were removed and analyzed by LC-MS using a linear gradient (water/MeCN 5%/95% → MeCN 100%, 0-8 min with 0.1% formic acid) and a (C-18 column (4.6 × 50 mm, 1.8 μm) (Agilent). The photoproducts of DEAC-OXM were generated and analyzed in an analogous manner. The integrals of the remaining caged molecule from each sample were normalized to the integral of the un-illuminated sample, averaged and plotted against time for the first two minutes of illumination. Linear regression provided a measurement of slope, and the ratio of the two slopes was used to determine the relative photolysis curve.

### *In vitro* GPCR activation assays

GloSensor assay of G-protein signaling. Human embryonic kidney 293T cells were grown in Complete DMEM (Dulbecco’s modified Eagle’s medium (Invitrogen, Carlsbad, CA) containing 5% fetal bovine serum (Corning), 50 U/mL Penicillin-Streptomycin (Invitrogen), and 1 mM sodium pyruvate (Corning)) and maintained at 37 °C in an atmosphere of 5% CO_2_ in 10 cm TC dishes. Media in 10 cm TC dishes with HEK 293T cells (at around 70% confluence) were replaced with Opti-MEM (Invitrogen). Then the SSF-MOR plasmid, GloSensor (Promega) 22F cAMP dependent reporter plasmid (Promega), and Lipofectamine 2000 (Invitrogen) in Opti-MEM were added. The dishes with transfection media were incubated at 37°C in an atmosphere of 5% CO_2_ for 6 h before replacing media with complete DMEM. After incubating at 37°C in an atmosphere of 5% CO_2_ for 16 h, transfected cells were plated in ploy-*D*-lysine coated 96-well plates at ∼40,000 cells/well and incubated at 37 °C in an atmosphere of 5% CO_2_ for 16 h. On the day of assay, media in each well were replaced with 50 µL of assay buffer (20 mM HEPES, 1x HBSS, pH 7.2, 2 g/L d-glucose), followed by addition of 25 µL of 4x drug solutions for 15 min at room temperature. To measure the activation of Gi-coupled opioid receptors, 25 µL of 4 mM luciferin supplemented with isoproterenol at a final concentration of 200 nM was added, and, following gentle mixing, luminescence counting was performed using a plate reader (iD5, Molecular Devices) after 25 min. For antagonism experiments, the candidate antagonist (naloxone, PhNX, PhOX) was added to the assay buffer (50 uL/well) at 2x the final concentration and allowed to incubate for 5 minutes prior to the addition of the MOR agonist DAMGO.

### Data analysis

MOR activation was expressed as % of DAMGO maximal effect and concentration-response curves were fitted using Prism 9 (Graphpad Software, La Jolla, CA, USA).

### β-arrestin recruitment assay

HEK-293T cells were seeded on 6-well culture plates at 3 x 10^6^ cells/well and grown in Dulbecco modified Eagle medium (DMEM; Thermo Fisher Scientific, Pittsburg, PA, United States) supplemented with L-Glutamine 200 mM, Sodium Pyruvate 100 mM and MEM non Essential Amino Acids 100X (Biowest). 10% fetal bovine serum (FBS; Merck KgaA, Darmstadt, Germany), streptomycin (100 μg/mL), and penicillin (100 μg/mL) in a controlled environment (37°C, 98% humidity, and 5% CO2). 24 h hours after seeding, cells were transfected the split NanoBiT® vectors NB MCS1 (Promega, Madison, WI, United States) fused to β-arrestin2 or the human MOR (0.1 µg β-arrestin2-LgBIT cDNA, 2 µg of the hMOR-SmBIT cDNA) using polyethylenimine (PEI; Polysciences EuropeGmbH, Hirschberg an der Bergstrasse, Germany) in a 1:3 DNA:PEI ratio. 48 h after transfection, cells were rinsed, harvested, and resuspended in 4 ml/well of Hanks’ Balanced Salt solution (HBSS, Sigma Aldrich, Switzerland). Cells (80 μl/well) were then plated in 96-well white plates (PO-204003, BIOGEN) and immediately treated with increasing concentrations of DAMGO or PhOX (0.1 nM to 10 µM), 5 minutes later at 2 µM elenterazine (Prolume Ltd) was added and luminescence (490 to 410 nm) was measured during 6 min using a CLARIOstar (BMG Labtech) plate reader.

### Data analysis

β-arrestin recruitment was expressed as % of DAMGO maximum effect and concentration-response curves were fitted using Prism 9 (Graphpad Software, La Jolla, CA, USA).

### *Ex vivo* occupancy

Mice with optical fiber implants in the anterior dorsal striatum were injected with vehicle or PhOX (30 mg/kg, *s.c.*). 30-min post-injection, photoactivation was performed using an Arduino-controlled 375nm laser (Vortran, 10 x 200 ms, 1 Hz, 30 mW). Brains were harvested 30-seconds post-laser stimulation and stored at -80 °C. Frozen tissue was sectioned (20 µM) on a cryostat (Leica, Germany) and thaw mounted onto Superfrost Plus glass slides (Avantor, USA). Brain slices were incubated 10 minutes in buffer (50 mM Tris-HCL, 10mM MgCl2) containing [3H]DAMGO (5 nM). Following incubation, slides were air dried and apposed to a BAS-TR2025 Phosphor Screen (Fujifilm) for 5-10 days and imaged using a phosphorimager (Typhoon FLA 7000).

### Data analysis

Quantification of 3H-DAMGO was performed in ImageJ (NIH). Images were calibrated using a C-14 reference to convert grey pixel values to nCi/g. ROIs were then drawn around the dorsal striatum of sections proximal to the fiber placement. Average mean measurements per mouse (n = 5 mice, 5-7 sections each) were plotted to compare ipsilateral (left) to contralateral (right) hemispheres.

### [^18^F]-Fluorodeoxyglucose PET

This procedure was based on previous studies (Cai *et al*., 2019; Bonaventura *et al*., 2021). One week before the PET procedure, surgeries to implant a fiber optic in the VTA were performed as described. Mice were habituated to experimenter handling, patch cord tethering, and the open field arena prior to experiment day. Mice were fasted 16 h before the experiment. On the day of the experiment, mice (n = 6-8 per condition) received a subcutaneous injection of vehicle (buffered saline), PhOX (30 mg/kg), or Oxymorphone (10 mg/kg). Thirty minutes after drug injection, mice were injected intraperitoneally with 0.35 mCi of 2-deoxy-2-[^18^F]fluoro-D-glucose (FDG; Cardinal Health) and placed into open-field chambers for drug photouncaging during uptake. Immediately after FDG administration single pulses of 375 nm light (200 ms, 30 mW) were delivered every 10 min (“LIGHT ON”), or no light was delivered (“LIGHT OFF”). After 30 min, mice were anesthetized with 1.5% isoflurane, placed on a custom-made bed of a nanoScan small animal PET/CT scanner (Mediso Medical Imaging Systems) and scanned for 20 min on a static acquisition protocol, followed by a CT scan.

### Data analysis

The PET data were reconstructed and corrected for deadtime and radioactive decay. (PMOD Technologies, Zurich, Switzerland). The PET data were coregistered to an MRI template (Ma *et al*., 2005) using the PMOD software environment (PMOD Technologies, Zurich, Switzerland). The coregistered PET data was analyzed in a voxel-based statistical analyses using a one-way ANOVA with four levels: vehicle LIGHT ON, Oxymorphone, PhOX LIGHT ON and PhOX LIGHT OFF. The values of the statistical parameter T for each contrast between conditions were mapped on the MRI template and filtered for clusters of voxels containing more than 100 contiguous voxels with statistical significance above the threshold (p < 0.05). All statistical parametric mapping analyses were performed using Matlab R2021a (Mathworks) and SPM12 (University College London). All qualitative and quantitative assessments of PET images were performed using PMOD. After the PET experiment animals were sacrificed and fiber placement was evaluated as described below, no animals were excluded from the analysis based on fiber placement.

### Brain slice preparation

Animal handling protocols were approved by the UC San Diego Institutional Animal Care and Use Committee. For synaptic transmission experiments, postnatal day 15−35 mice of both sexes on a C57/Bl6 background were used while GIRK experiments were performed on mice of the PValbCre/Ai14 background. Mice were sacrificed by gas anesthetic followed by rapid decapitation. Brains were removed in ice-cold choline solution equilibrated with 95% O2/5% CO2 (carbogen) containing (in mM) 25 NaHCO3, 1.25 NaH2PO4, 2.5 KCl, 7 MgCl2, 25 glucose, 0.5 CaCl2, 110 choline chloride, 11.6 ascorbic acid, and 3.1 pyruvic acid, and chilled for 3 minutes. Brains were then blocked to generate sections from the horizontal plane and mounted in a VT1000S vibratome (Leica Instruments). Slices (300 μm) were cut in ice-cold choline solution perfused with carbogen. Slices were recovered for 30 minutes at 34 °C in a holding chamber containing carbogenated artificial cerebrospinal fluid (ACSF) composed of (in mM) 125 NaCl, 2.5 KCl, 25 NaHCO3, 1.25 NaH2PO4, 2 CaCl2, 1 MgCl2, and 15 glucose, osmolarity 295 (note 2.7 g glucose used for these experiments), after which the holding chamber was moved to room temperature until use in experiments.

### Electrophysiology

Recordings were performed on slices in a submerged slice chamber perfused with 32 °C ACSF equilibrated with 95% O2/5% CO2. Whole-cell voltage clamp recordings were made with an Axopatch 700B amplifier (Axon Instruments). Data were filtered at 3 kHz, sampled at 10 kHz, and acquired using National Instruments acquisition boards and a custom version of ScanImage written in MATLAB (Mathworks). Cells were rejected if holding currents exceeded −200 pA or if the series resistance (<25 MΩ) changed during the experiment by more than 20%.

All recordings were performed within 5 hours of slice cutting in a submerged slice chamber perfused with ACSF warmed to 32°C and equilibrated with carbogen. The hippocampal CA1 region was identified by morphology. Pyramidal cells were patched with pipets (open pipet resistance 2.8-3.5 MΩ) containing cesium low chloride internal solution composed of (in mM) 135 CsMeSO3, 3.3 QX314-Cl, 10 HEPES, 4 MgATP, 0.3 NaGTP, 8 Na2-phosphocreatine, 1 EGTA (CsOH). For synaptic transmission experiments, excitatory transmission was blocked by the addition of CPP (10 µM) and NBQX (10 µM) to the bath. Once the whole cell configuration was obtained, pyramidal cells were voltage clamped at 0 mV to facilitate detection of outward currents. IPSCs were electrically evoked (two pulses, 0.5 ms, 50-300 µA, 50 ms interval) every 20 seconds by a bipolar theta glass stimulating electrode (∼5 µm tip diameter) placed at the border between stratum pyramidale and stratum oriens approximately 50-150 µm from the patched cell. After establishing a stable baseline for 3-5 minutes, drugs were added to the bath at the times and concentrations indicated in their corresponding figure.

For analysis of GIRK currents, parvalbumin interneurons within the CA1 were identified by reporter expression in PVCre; TdTomato mice. PV+ cells were voltage clamped at -55 mV and GIRK currents were isolated by the addition of CPP (10 µM), NBQX (10 µM), picrotoxin (10 µM), and tetrodotoxin (1 uM) to the bath. A potassium based internal was used that contained (in mM) 135 KMeSO3, 5 KCl, 5 HEPES, 4 MgATP, 0.3 NaGTP, 10 phospho-creatine, 1.1 EGTA (KOH).

In both experiments, after recordings stabilized, a 50 ms flash of UV light was applied from the 365 nm-UV channel of a pE-300^white^ LED (CoolLED) reflected through a 60× LUMPLANFL 1.0 NA objective (Olympus) on SliceScope Pro 6000 microscope (Scientifica) with a 405 nm long-pass dichroic mirror (Di02-R405-25x36, Semrock) mounted in the fluorescence turret. Light power was set to 5 mW in the sample plane (∼120 mW of an ∼20 mm diameter “beam” at the back aperture). The resulting illumination field was ∼0.02 mm^2^.

### Data analysis

Data from electrophysiology experiments were analyzed in Igor Pro (version 6.02A, Wavemetrics). For synaptic transmission experiments, IPSCs were calculated by averaging a 2 ms window around the peak, and normalized to the average of the three sweeps (one minute) prior to uncaging or drug addition. To determine percent suppression, the average IPSC amplitude from minutes 2-3 post uncaging flash was compared to the minute prior to uncaging. The paired pulse ratio (PPR) was determined by Peak 2/Peak1. For GIRK analysis, peak drug effect was determined by comparing the average of three sweeps (15 seconds) to the minute prior to drug addition. For determination of uncaging effect, the 4 seconds prior to the uncaging flash were used as the baseline, and the current at seconds 5-10 post uncaging were analyzed to calculate suppression. Time constants were calculated by a mono-exponential fit to time 0-10 seconds post uncaging.

### Blood-brain barrier penetrance and pharmacokinetics

Pharmacological studies were conducted at the UCSD Translational Pharmacology and Bioanalysis Laboratory. For blood-brain barrier studies, mice were anesthetized briefly with isoflurane and injected intravenously (retro-orbital) with either PhOX (8.5 mg/kg), Oxymorphone freebase (5 mg/kg), PhNX (15 mg/kg) or NLX HCl (10 mg/kg). 15 minutes post-injection, mice were anesthetized again and a retro-orbital heparinized blood sample was removed (55-65 µl), followed by cardiac perfusion with room temperature PBS (20-25 mL).

Brains were then collected, immediately frozen on dry ice, and stored at -80C. Subsequently the brain tissue was homogenized through a series of extraction-filtration procedures. For half-life determination, mice were injected *i.p.* with PhOX and retro-orbital blood samples were taken at the following time points: 5 minutes, 15 minutes, 1 hour, and 2 hours post-dose. Plasma samples were extracted from blood samples via centrifugation at 5,000 rpm for 5 minutes.

### Data analysis

Tissue and plasma concentrations were determined using LC-MS/MS. Standards were used to generate an external calibration curve in blank plasma or tissue homogenate using a linear regression algorithm to plot the peak area ratio vs concentration with 1/x weighting, over the full dynamic range of analyte concentrations.

### Behavior

### Tail-flick

Tail-flick analgesia experiments were conducted using a TF-2 Tail-Flick Apparatus (Columbus Instruments). Mice implanted with optical fibers in the vlPAG or VTA were pre-handled and then manually restrained during the experiment. Mice were tested twice on two consecutive days (with and without NLX); the conditions were counterbalanced across days. 15 minutes after injection of PhOX (15 mg/kg, *i.v.*) or PhOX + NLX (15 mg/kg, 10 mg/kg, *i.v.*), the tail was positioned flat 15 mm from 6 W heat beam and latency to tail flick or time out (30 s) was measured. A baseline measurement was taken at 25 min after drug injection and multiple measurements were taken after application of a train of 375 nm light flashes (10 x 200 ms, 1 Hz, 30 mW). The experimenter was blind to drug treatment conditions.

### Hargreaves with capsaicin hyperalgesia

Hargreaves analgesia experiments were performed mice implanted with optical fibers in their left NAc-mSh. Mice were injected with PhOX (12 mg/kg, *s.c.*) and set on a hargreaves heated glass platform (32°C) to acclimate for 17 minutes. The Hargreaves apparatus (Model 400, IITC life sciences) heat source was set to 50% threshold with a cutoff time of 20 seconds. Heat was applied to the right contralateral paw for all latency measurements. Paw withdrawal latencies were characterized as a sudden, responsive lifting, licking, or shaking of paw and measurements were not taken when the subject was moving or grooming. To induce, capsaicin hyperalgesia for thermal nociception (Rashid *et al*., 2003) a pharmaceutical grade 0.1% capsaicin cream was applied to the hindpaw. A pre-capsaicin baseline was measured at 17 minutes post-injection.

Capsaicin was lightly applied to the hindpaw at 20 minutes post-injection, a post capsaicin baseline was taken at 28 minutes post-injection, photolysis (10 x 200 ms, 1 Hz, 45 mW) occurred at 29 minutes post-injection of PhOX. The first post-uncaging latency was 1 minute after photolysis, subsequent latencies were taken at 5, 10, 20, and 30 minutes. In the no light control experiment, mice were fiber tethered with all other parameters identical except omission of photolysis. The 375 nm laser was coupled to a commutator to allow mice freedom of movement.

### Hot plate

Mice were injected with either saline or PhOX (15 mg/kg, *i.v.*) and tested on the hot plate 15 minutes later. The average latency of 3 individual trials (1 every 5 min) is reported for each condition. Mice were next injected with morphine (5 mg/kg, *s.c.*) and 30 minutes after injection mice were again tested on the hot plate 3 times.

### Locomotion

For open field experiments with intravenous PhOX, mice on a on a C57Bl/6J background were implanted unilaterally on the VTA and allowed to recover. All mice were handled and acclimated to tethering to the fiber optic cable prior to photoactivation experiments. For intravenous (*i.v.*) injections, mice were briefly anaesthetized with 3-5% isoflurane and given retro-orbital injections of either PhOX (15 mg/kg) or PhOX + naloxone (15 mg/kg + 10 mg/kg) or PhOX + PhNX (15 mg/kg + 30mg/kg). After recovery from anesthesia, mice were tethered to an optical fiber(200 µm, 0.22 NA) coupled to a commutator connected to an Arduino-controlled 375 nm laser (Vortran). They were placed into a square enclosure (18 x 18 cm) where their position was video recorded at a 30 fps using a webcam (Logitech) fed into Smart 3.0 video tracking software (Panlab). After a 15-minute baseline, PhOX was photoactivated in the VTA (30mW power, 200ms, 1Hz, 10 flashes). Locomotion was recorded 15 minutes post stimulation.

For open field experiments comparing the locomotor response to subcutaneous PhOX after 15 or 30 minutes post-injection, PhOX (30 mg/kg *s.c.*) was injected and VTA photoactivation with a 375 nm laser (10 x 200ms, 1 Hz, 30mW) was conducted either 15 minutes or 30 minutes post-injection. For open field experiments comparing doses of subcutaneous PhOX, the following dose of PhOX were injected: 2.5 mg/kg, 5 mg/kg or 10 mg/kg *s.c.* 15 minutes post-injection, photoactivation in the VTA was conducted (10 x 200 ms, 1 Hz, 30 mW) and mice were recorded for 15 minutes post photoactivation. For open field experiments comparing the locomotor response to VTA PhOX at 3 vs. 10 light flashes, mice were injected with PhOX (10 mg/kg, *s.c.*) and placed into an open field. 15 minutes post-injection, photoactivation in the VTA was conducted (3 or 10 x 200 ms, 1 Hz, 30 mW).

For comparison of LED and UV locomotor behavior after VTA PhOX activation, mice were injected with PhOX (30 mg/kg, *s.c.*) and after 10 minutes connected to light sources set to deliver the following light powers for photoactivation: 375 nm laser (15 mW), 385 nm LED (2.5 mW), or 365 nm LED (1.4 mW). The reported power levels in this experiment were measured from the tip of a ferrule-connected optical fiber similar to those used for the implants. Optical fibers connected to mice were coupled directly to light sources without the use of a commutator in these experiments to maximize LED light power output. Mice were placed into the open field and locomotor behavior was recorded using Smart 3.0 (Panlab) video tracking software. After an initial baseline of 20 minutes photolysis protocol was initiated (10 x 200 ms, 1 Hz, one bout every 5 min) and locomotion post-photoactivation was recorded for 20 minutes.

### Data analysis

Velocity and total distance traveled was quantified using Smart 3.0 video automated tracking software (Panlab). Velocity (cm/s) was calculated by quantifying distance traveled in time bins of 5 seconds and dividing by seconds.

### Conditioned place preference

For conditioned place preference experiments, a custom built 2-chamber apparatus (26 x 26 cm, UCSD Machine Shop) was used with both visual and tactile cues. Before conditioning, mice were habituated for 20 minutes by being allowed to freely roam both chambers. Pre-preference was recorded and PhOX + light was paired with the least preferred chamber. Prior to conditioning sessions, mice were injected with PhOX (15 mg/kg, i*.v.)* or vehicle (saline, *i.v.*) under anesthesia and allowed to recover for 15 minutes. For conditioning without light stimulation: Conditioning took place over 4 days, where PhOX injection was paired with one context and vehicle was paired with the other on alternating days. Conditioning sessions took place over 20 minutes. On test day, mice were allowed to roam both chambers for 20 minutes. For conditioning with light stimulation in the VTA, mice were implanted with 200 μm optical fibers as previously described.

During conditioning sessions mice were tethered to a fiber optic cable attached to a 375 nm laser via a commutator and exposed to a uncaging stimulus (10 x 200 ms, 1Hz) every 5 minutes. For reversal conditioning with VTA stimulation, PhOX plus light stimulation was paired with the previously least preferred chamber from conditioning test day of Phase 1 and PhOX plus naloxone (with light) was paired with the previously most preferred chamber.

### Data analysis

Place preference was quantified as time spent in each paired chamber using Smart3.0 (Panlab) automated tracking software.

### Two-site fiber photometry and uncaging

For fiber photometry with dLight1.3b, mice were injected with dLight1.3b +mCherry in the left NAc-mSh and after at least 3 weeks of expression, they were implanted with 200 μm optical fibers into the left NAc-mSh and left VTA. Mice were allowed to recover at least one week post fiber implantation. Fiber photometry recordings were done using a commercially available fiber photometry system FP3002 (Neurophotometrics). For dLight1.3b experiments with PhOX, mice were injected with either PhOX (15 mg/kg, *i.v.*) or PhOX + NLX (15 mg/kg, 10 mg/kg, i*.v.*). 5 minutes after injection, mice were connected to two fiber optic cables, one coupling the NAc-mSh optical fiber implant to the fiber photometry system (470 nm LED for dLight1.3b and 560 nm LED, mCherry control wavelength) and the other coupling the VTA optical fiber to a 375 nm Arduino-controlled laser (Vortran. After a 10-minute baseline, photoactivation in the VTA was applied with a single 375 nm light pulse (200 ms, 50 mW). dLight1.3b fluorescence was recorded 20 minutes post photoactivation. For fiber photometry experiments with dLight1.3b testing for the effects of off-target wavelength photostimulation in the VTA, fiber photometry with PhOX was performed as described using a 473 nm laser (Optoengine) instead of a 375 nm laser. Fiber photometry experiments without PhOX injection were performed as described except no PhOX injection was administered. Experiments examining the power-response relationship between VTA PhOX photactivation and NAc-mSh dLight1.3b were conducted on separate days as described for the following light powers: 1 mW, 5 mW, 20 mW, and 50 mW.

For repeated uncaging with VTA PhOX and NAc-mSh dLight1.3b, mice were injected with PhOX (10 mg/kg, *s.c.*) and after 30 minutes post-injection a 375 nm light pulse (20 mW, 200 ms) was delivered to the VTA. After 10 minutes, a 2^nd^ light pulse was delivered. dLight1.3b signal was recorded 10 minutes post each uncaging event.

For site-separated fiber photometry with PhNX, mice were injected with dLight1.3b and mCherry (control) in the NAc-mSh and implanted with optical fibers in the NAc-mSh and VTA. Mice were anaesthetized and injected with PhNX (30 mg/kg, *i.v.*). After recovery, mice were connected to two fiber optic cables, one coupling the NAc-mSh optical fiber implant to the fiber photometry system (470 nm LED for dLight1.3b and 560 nm LED, mCherry control wavelength) and the other coupling the VTA optical fiber to a 375 nm Arduino-controlled laser. Baseline fluorescence was recorded for 10 minutes. Photoactivation in the VTA (20 x 200 ms x 80 mW, 1 Hz) was applied 1 minute prior to morphine (10 mg/kg, *i.p.*) injection, and two times every 5 minutes after morphine injection. Post-photoactivation, fluorescence was recorded for 30 minutes. Mice receiving morphine only were anaesthetized and allowed to recover with no injection. Mice were tethered to optical fiber cables attached to the fiber photometry system and 375 nm cables as described previously. A 10-minute baseline was recorded and then morphine (10 mg/kg, *i.p.*) was injected. Afterwards, 30 minutes of fluorescence was recorded.

### Fiber photometry with chronic morphine

For chronic morphine experiments, mice were injected with GRAB-DA2m (Sun *et al*., 2020) in the left NAc-mSh and implanted with optical fibers in the left NAc-mSh and left VTA. Prior to chronic morphine treatment, mice were handled, habituated to tethering, and their baseline response to PhOX were tested. For PhOX photoactivation in the VTA, PhOX (12 mg/kg, *s.c.*) was injected 30 minutes prior to laser stimulation. A 5 minute baseline of GRAB-DA2m signal in the NAc-mSh was recorded prior to photostimulation (10 x 200 ms, 1 Hz, 20 mW). For chronic morphine treatment, mice received two daily injections of morphine or saline (11:00 am and 7:00 pm) using an intermittent escalating dose protocol (Taylor *et al*., 2016), of 10, 20, 30, 40 and 50 mg/kg *i.p.* morphine. On day 5, fiber photometry recording in the NAc-mSh of GRAB-DA mice with PhOX uncaging was performed as described above in the morning and mice received their 2 daily morphine injections afterwards. Following 4 days of abstinence, fiber photometry with PhOX uncaging was performed again.

### Data analysis

Data analysis including peri-event time histograms of ΔF/F and z-score values surrounding photoactivation events, AUC, and tau calculations were conducted using Matlab R2020a (Mathworks) and custom scripts in Igor Pro (Wavemetrics). Analysis in Matlab was conducted using pMAT an open source fiber photometry analysis package (Bruno *et al*., 2021). The signal (470 nm) and control (560 nm) channel were individually smoothed and downsampled and bleach corrected. A scaled control channel was calculated using a least-squares regression to normalize the scale of the channels. ΔF/F values were calculated using the following equation: ΔF/F = (Signal Channel-Scaled Control Channel)/Scaled Control Channel. A robust z-score was generated using the following formula: [ΔF/F Event – median (ΔF/F baseline)]/ median absolute deviation (MAD) of baseline.

### Same-site fiber photometry and uncaging

For same-site fiber photometry with PhOX, vGAT-Cre mice were transduced with cre-dependent GCaMP8s in the left RMTg and implanted with an optical fiber implanted over the left VTA. Mice were tethered to an optical fiber attached to a custom-built optical relay (Supporting Figure 11) that was connected to both the fiber photometry system and a 375 nm laser (Oxxius). Prior to PhOX injection, laser stimulation was applied to the VTA to control for the effect of light delivery alone on fluorescence signal. A 10-minute baseline was recorded, then a single, 20 mW, 200 ms laser pulse was delivered every 10 minutes for a total of 3 flashes. After the preinjection test session, PhOX (30 mg/kg, *s.c.*) was injected while mice were still tethered to the fiber optic cable. 20 minutes after injection, Fiber Photometry recording was resumed. A 10 minute baseline was recorded, and then a 200 ms, 20 mW flash was delivered, and 20 minutes was recorded post-flash.

### Data analysis

Same-site fiber photometry analysis was conducted as described for site-separated fiber photometry with the addition of laser light artifact correction. For light-artifact correction, the average ΔF/F of 3 preinjection flashes was subtracted from the post-injection ΔF/F.

### Surgeries

For behavior, FDG PET or binding assay experiments, 200 µm optical fibers were implanted at the following coordinates (in mm, from bregma), for unilaterally in the left VTA at AP −3.3; ML −0.5 with a 10° angle; DV −4.15, bilaterally in the PAG at AP -4.60, ML ±0.32; DV -2.75 with a 10° angle, left NAc-mSh at AP 1.54, ML -0.48, and DV -4.28, or left aDS AP: 1.2; ML: -1.4; DV: -3.00. Implants were secured with light-cured dental cement. For site-separated fiber photometry experiments: 400nl of AAV1-hsyn1-dLight1.3b (Addgene) or AAV9-CAG-dLight1.3b (Addgene) or AAV9-hsyn-GRAB-DA2m(DA4.4) (WZBiosciences) mixed 1:9 with AAV5-hSyn-mCherry were injected into the left NAc-mSh at AP 1.54 mm from bregma;ML 0.48 mm; DV - 4.28 mm. After at least 3 weeks post viral injection mice were implanted with 200 µm optical fibers. Left NAc-msh implants were fluorescence guided at approximately 0.3 mm above viral coordinates and left VTA implants were at AP −3.3 mm from bregma; ML −0.5 mm with a 10° angle; DV −4.15 mm or left PAG implants at AP -4.60 mm from bregma, ML 0.32 mm; DV -2.75 mm with a 10° angle. For same-site fiber photometry experiments 400 nl pGP-AAV-syn-FLEX-jGCaMP8s-WPRE (AAV1) (Addgene) mixed 1:9 with AAV5-hSyn-mCherry was injected into left RMTg of vGAT-cre mice at AP -3.9 mm from bregma; ML –0.4 mm; DV 4.5 mm. After allowing for at least 3 weeks of expression, 200 µm optical fibers were implanted into the left VTA using fluorescence guidance, aiming for at AP −3.3 mm from bregma; ML −0.5 mm with a 10° angle; DV −4.15 mm. All mice were given at least one week to recover after fiber implant surgeries.

### Histology

Mice were perfused using PBS followed by 4% paraformaldehyde and brains were collected, and switched to 30% sucrose prior to cryosectioning. 40 uM sections were taken and mounted with DAPI. For sections of the VTA, tyrosine hydroxylase immunohistochemistry (TH IHC) was carried as followed: free-floating brain slices were blocked in 5% normal donkey serum (NDS) in PBS plus 0.1% triton (PBST) for 2 hours and then transferred to 1:1000 TH anti-rabbit primary antibody (Millipore) with 1% NDS in PBST overnight at 4 °C, on day 2, the slices were washed 3 x 10 minutes in PBS at RT and transferred to second antibody Alexa 647 donkey anti-rabbit 1:1000 with 1% NDS in PBST for 1.5 hours at RT. After incubation with secondary antibody, slices were washed in 3 x 10 min in PBS RT and mounted with DAPI. Slices were imaged using a BZ-X100 fluorescence microscope (Keyence) and images were processed using ImageJ.

### Quantification and Statistical Analysis

For all comparisons in this manuscript, we describe the number of n’s, what the n represents, the statistical test used, and the definition of error bars in the figure legend. For every comparison related to a figure, the details of the statistics are in the figure legend. For every other comparison, they are listed in the results section. Comparisons were considered to have reached statistical significance if the p-value was less than 0.05, unless otherwise stated. The p-values that correspond to asterisks are listed in the figure legends. When appropriate, exclusion criteria are listed in the corresponding methods details section.

D’Agostino & Pearson and/or Shapiro-Wilk tests were used to determine whether data were normally distributed. For comparison between two groups, either an unpaired or paired two-sided t-test was used for normally distributed datasets, and either a Mann-Whitney U-test or Wilcoxon matched-pairs signed rank test was used for non-normally distributed data. For comparison between two or more groups, differences were detected using either a One-way ANOVA or a Friedman test, and interactions were identified using Two-way ANOVA. A post-hoc comparisons were used to determine differences between specific conditions (Tukey’s, Bonferroni’s, Sidak’s or Dunn’s). The source data spreadsheet available in the supporting information include the details of all statistical tests used in each figure.

## Supplemental Figures

**Figure S1.**
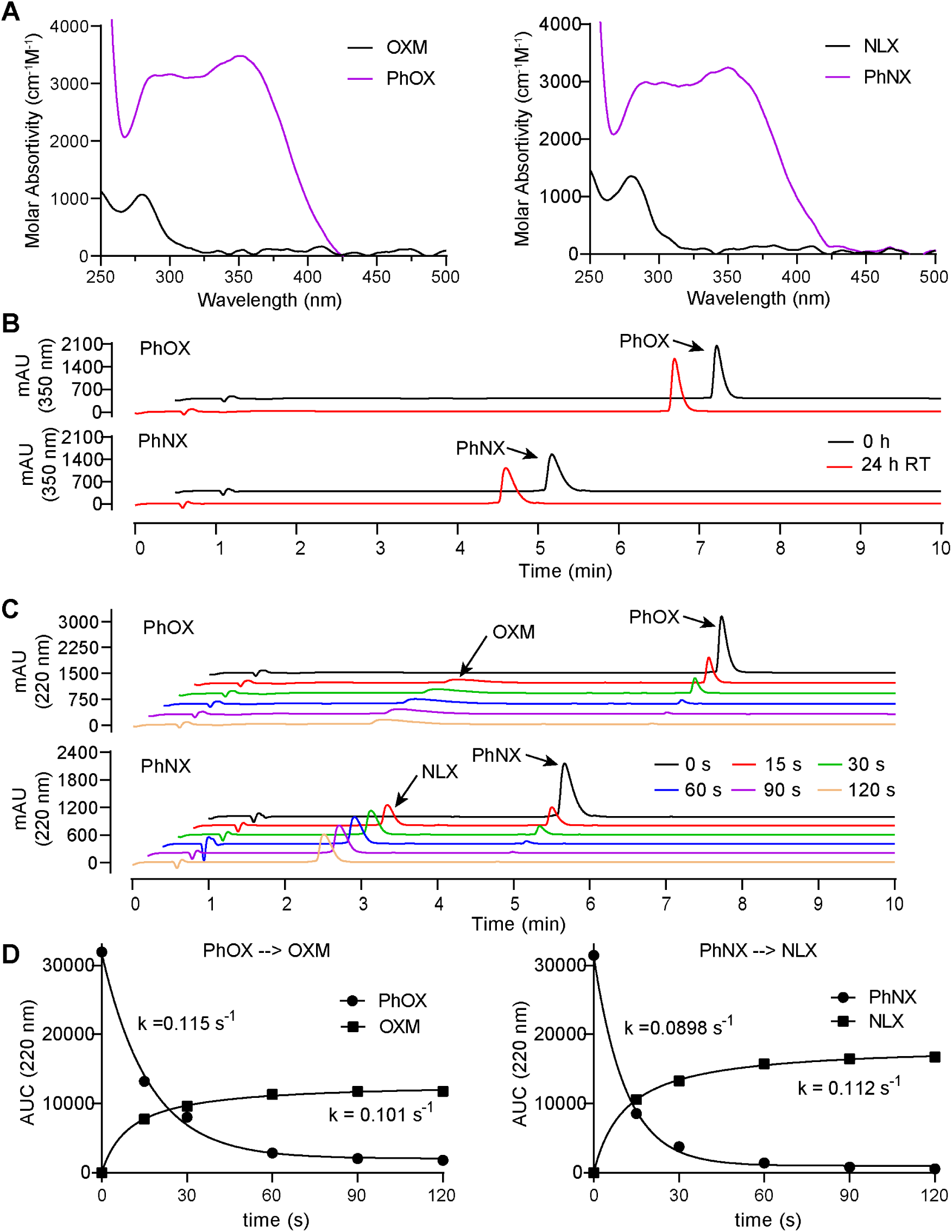
*In vitro* characterization of PhOX and PhNX stability and photolysis. **(A)** UV/VIS absorbance spectra of PhOX and OXM (left), and PhNX and NLX (right). The DMNPE caging group exhibits minimal absorbance beyond 425 nm, rendering it compatible with fluorescence imaging with visible light-excited fluorophores. **(B)** High pressure liquid chromatography (HPLC) chromatograms (350 nm absorbance) of samples of PhOX (top) and PhNX (bottom) (1 mM) dissolved in PBS buffer (pH 7.2) before and after being left in the dark at room temperature for 24 hours. The lack of change in the chromatograms indicates that both compounds are highly stable in the absence of illumination. **(C)** Waterfall plots of the HPLC chromatograms (220 nm absorbance) for samples of PhOX (top) and PhNX (bottom) (1 mM) dissolved in PBS buffer (pH 7.2) after being illuminated with 375 nm light for the indicated time periods. **(D)** Summary of PhOX (left) and PhNX (right) photouncaging reactions over time, as measured by HPLC (220 nm absorbance). The area under the curve (AUC) data include the examples shown in Supporting Figure 1C (n=3 samples per compound). The rate constants of starting material loss and product formation were obtained by fitting the curves with a single exponential function. All data are plotted as mean ± SEM.

**Figure S2.**
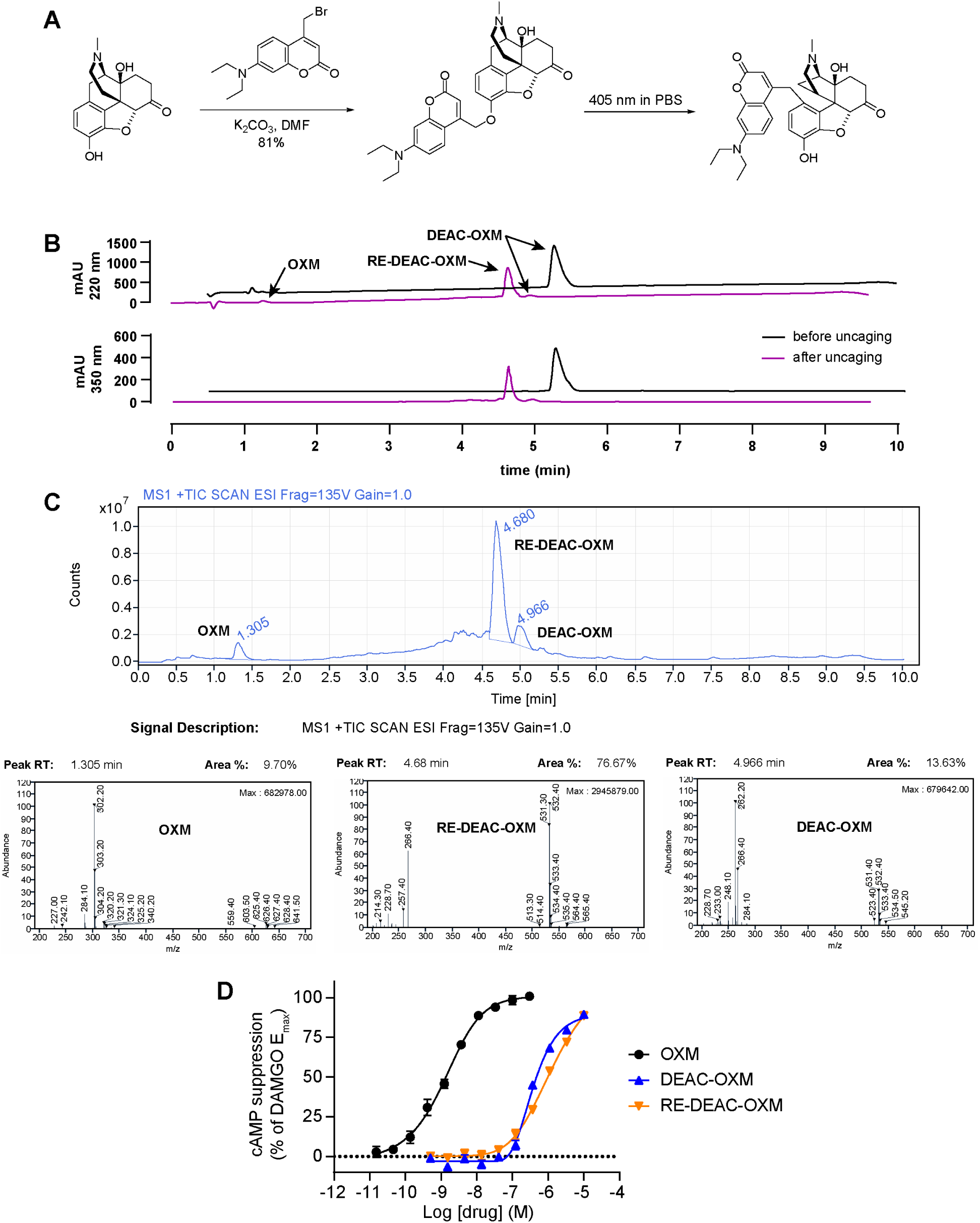
Synthesis and in vitro evaluation of DEAC-OXM. **(A)** Reaction scheme depicting the one-step alkylation procedure used to synthesize DEAC-OXM from commercially available oxymorphone (**2**) and DEAC-Br, as well as its photochemical conversion to the proposed primary reaction product “rearranged DEAC-OXM” (RE-DEAC-OXM), which likely occurs via a 1,4-Photo-Claisen rearrangement. **(B)** High pressure liquid chromatography (HPLC) chromatograms measured at 220 nm indicating predominant photoconversion of DEAC-OXM to RE-DEAC-OXM, which has a similar retention time, along with a much smaller amount of OXM in PBS (pH 7.2). **(C)** (Top) LC-MS (mass spectrometry) chromatogram of the reaction mixture shown in B. (Bottom) MS traces revealing that RE-DEAC-OXM has the same molecular weight as DEAC-OXM (531 Da). **(D)** Agonist dose-response curves at the MOR for DEAC-OXM, RE-DEAC-OXM, and OXM using the GloSensor™ assay of cAMP signaling in HEK293T cells. The solid line depicts the best-fit sigmoidal function used to derive the indicated EC_50_ value. OXM EC_50_ = 1.3 nM, DEAC-OXM EC_50_ = 380 nM, RE-DEAC-OXM EC_50_ = 1.3 µM. Data were normalized to the response produced by DAMGO (1 µM) and are expressed as mean ± SEM (n=5-10 wells per concentration).

**Figure S3.**
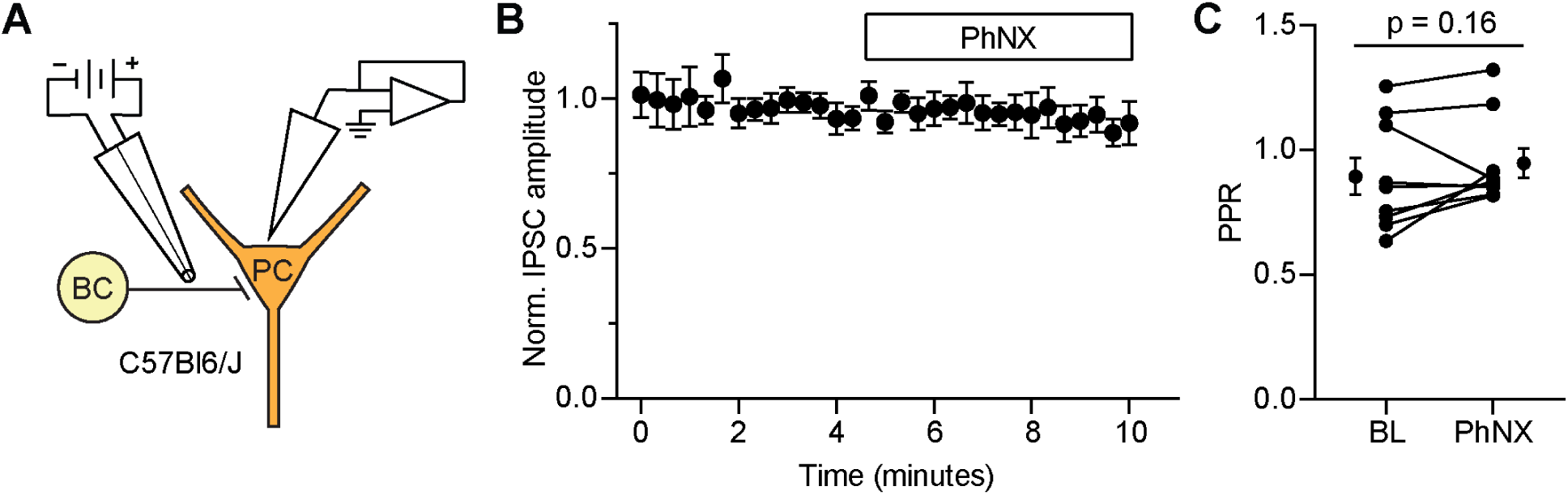
PhNX does not affect inhibitory synaptic transmission. **(A)** Schematic depicting whole-cell voltage clamp recording of opioid-sensitive synaptic inhibition in the CA1 region of hippocampus (same as Figure 3A). A small bipolar stimulating electrode constructed from theta glass was placed near the recorded pyramidal cell (PC) to preferentially stimulate basket cell (BC) axons. **(B)** Baseline-normalized inhibitory post-synaptic currents (IPSCs) in response to bath application of PhNX (1 µM), at the time indicated by the black line (n=9 cells from 7 mice). **(C)** Summary data of the pair-pulse ration (PPR) measured before and after PhNX application (n=9 cells from 4 mice, Wilcoxon signed rank). All data are plotted as mean ± SEM.

**Figure S4.**
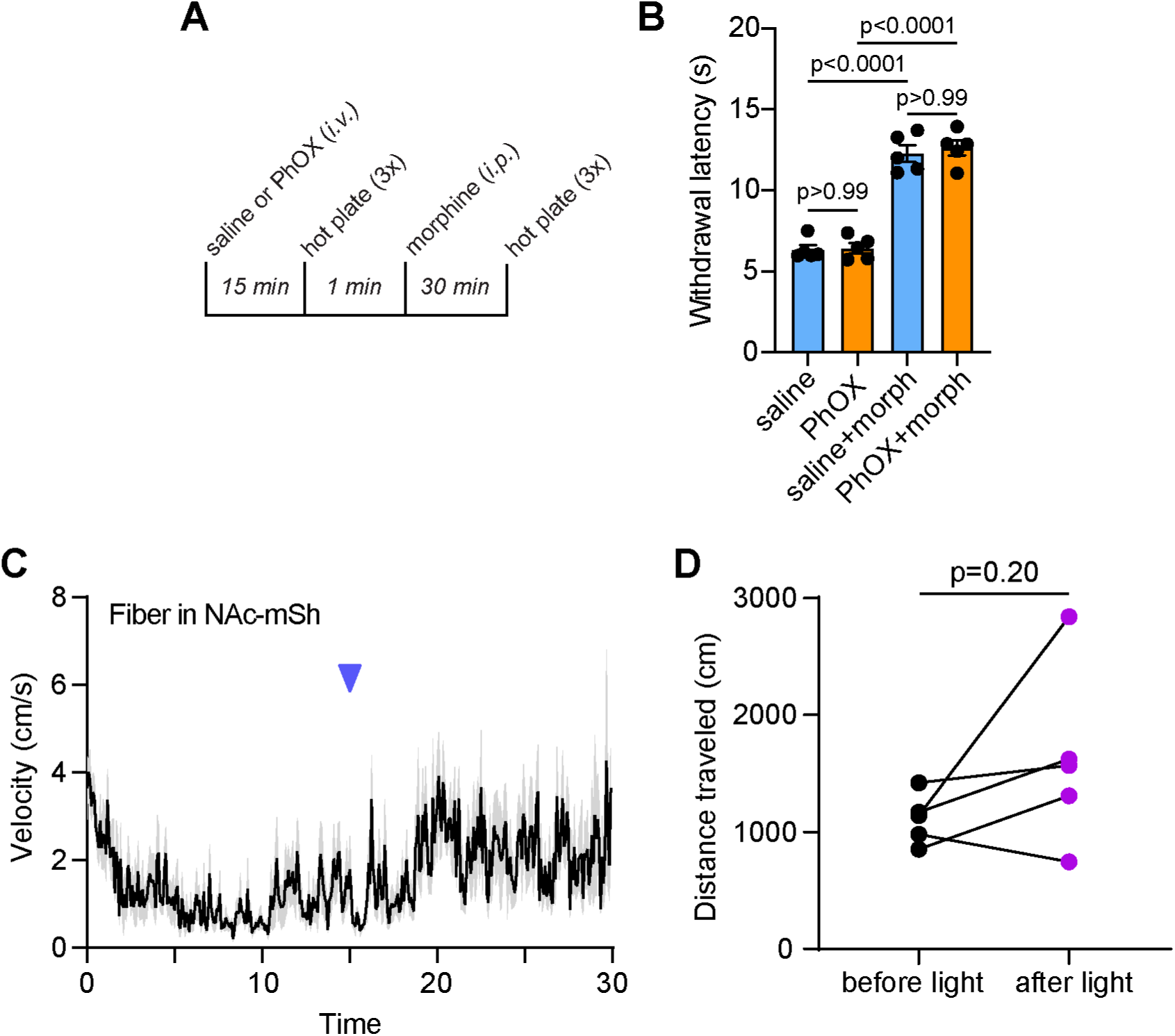
PhOX does not produce analgesia or alter systemic morphine analgesia in the absence of illumination. **(A)** Experimental timeline. **(B)** Hot plate paw withdrawal latencies observed after injecting mice with either saline or PhOX (15 mg/kg *i.v.*), followed by morphine (5 mg/kg, *s.c.*) (n=5 mice, One-way ANOVA, F(3,16)=73.21, p<0.0001, Bonferroni’s multiple comparisons test). **(C)** Average plot of velocity over time (n=5 mice) before and after PhOX photoactivation in the NAc-mSh. **(D)** Summary plot comparing the distance traveled in the 15 minutes before and after PhOX photoactivation (paired two-tailed t-test). All data are plotted as mean ± SEM.

**Figure S5.**
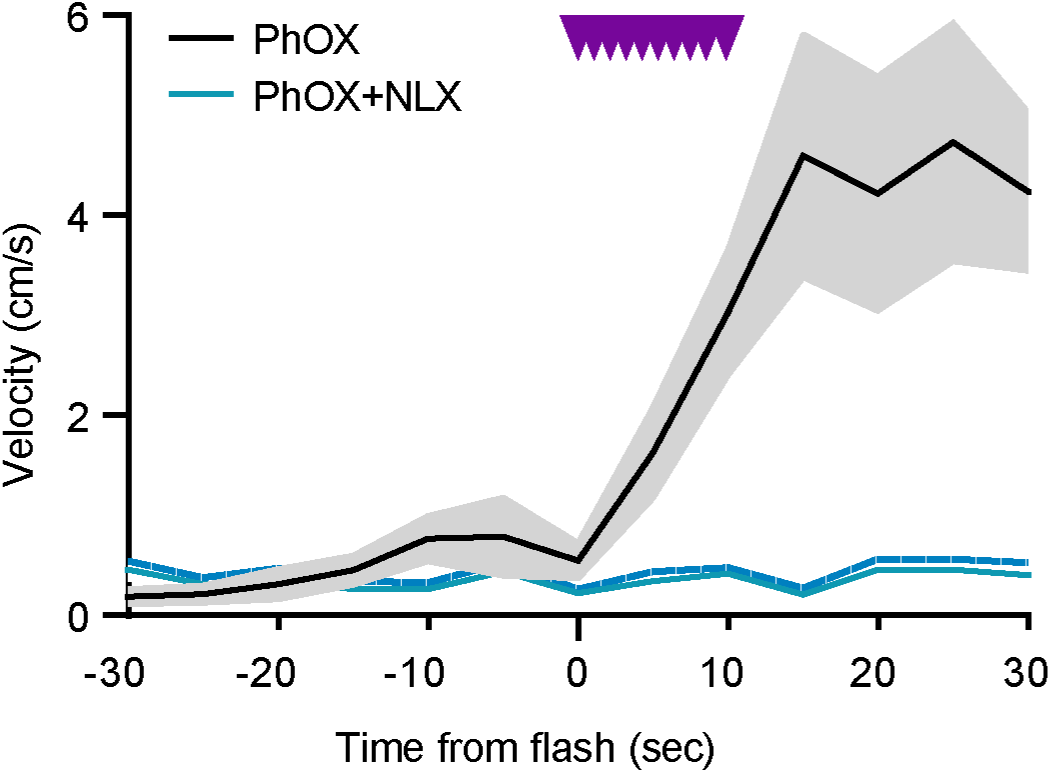
Unilateral PhOX photoactivation in the VTA drives locomotor activation within seconds of illumination. Data presented in Figure 6D zoomed in around the train of 375 nm light flashes. Data are plotted as mean ± SEM.

**Figure S6.**
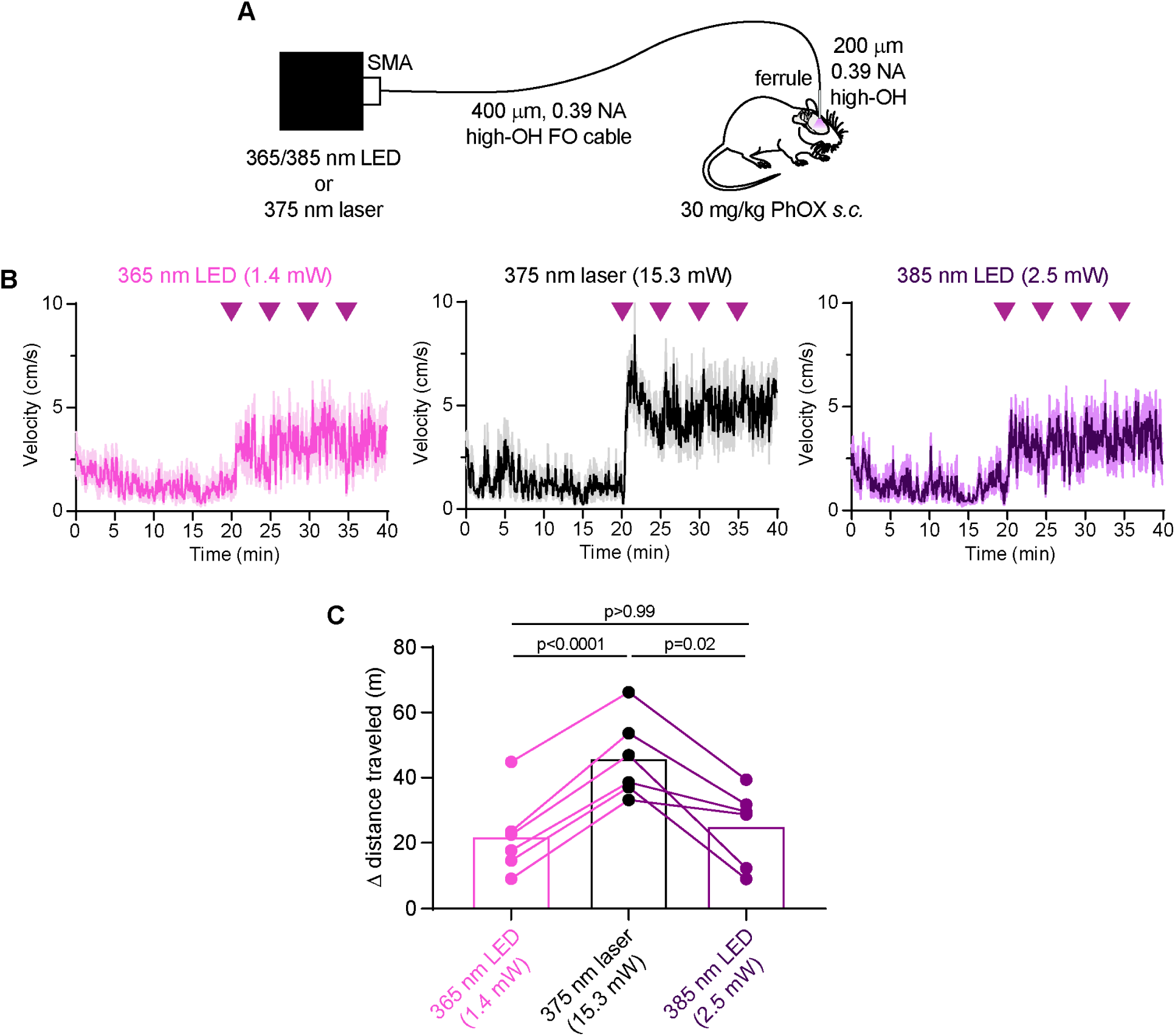
Comparison of locomotor responses evoked by PhOX photoactivation using low-cost LEDs. **(A)** Schematic indicating the direct coupling of light from either a 365 nm LED, a 385 nm LED, or a 375 nm laser into an optical fiber implanted over the VTA. The 400 µm, 0.39 NA high-OH coupling fiber was found to maximize the transfer of LED output to the output of a ferrule-coupled 200 µm, 0.39 NA optical fiber similar to those implanted in the mice. Maximal output from the 365 nm LED was 1.4 mW and 2.5 mW from the 385 nm LED. The 375 nm laser output was set to 15.3 mW in order to evoke a relatively strong locomotor response. **(B)** Average plots of velocity over time in the open field using trains of 10x200 ms flashes of light (2 Hz) at the times indicated by each arrow, using the indicated light sources (n=6 mice per condition). **(C)** Summary plot of the change in total distance traveled in the 20 minutes before and after light delivery (n=6 mice, Repeated measures one-way ANOVA, F(1.12,5.62)=20.99, p=0.004, Bonferroni’s multiple comparisons test). Neither LED at maximal power was as effective as 15.3 mW from the 375 nm laser, but both LEDs evoked similar locomotor responses. All data are plotted as mean ± SEM.

**Figure S7.**
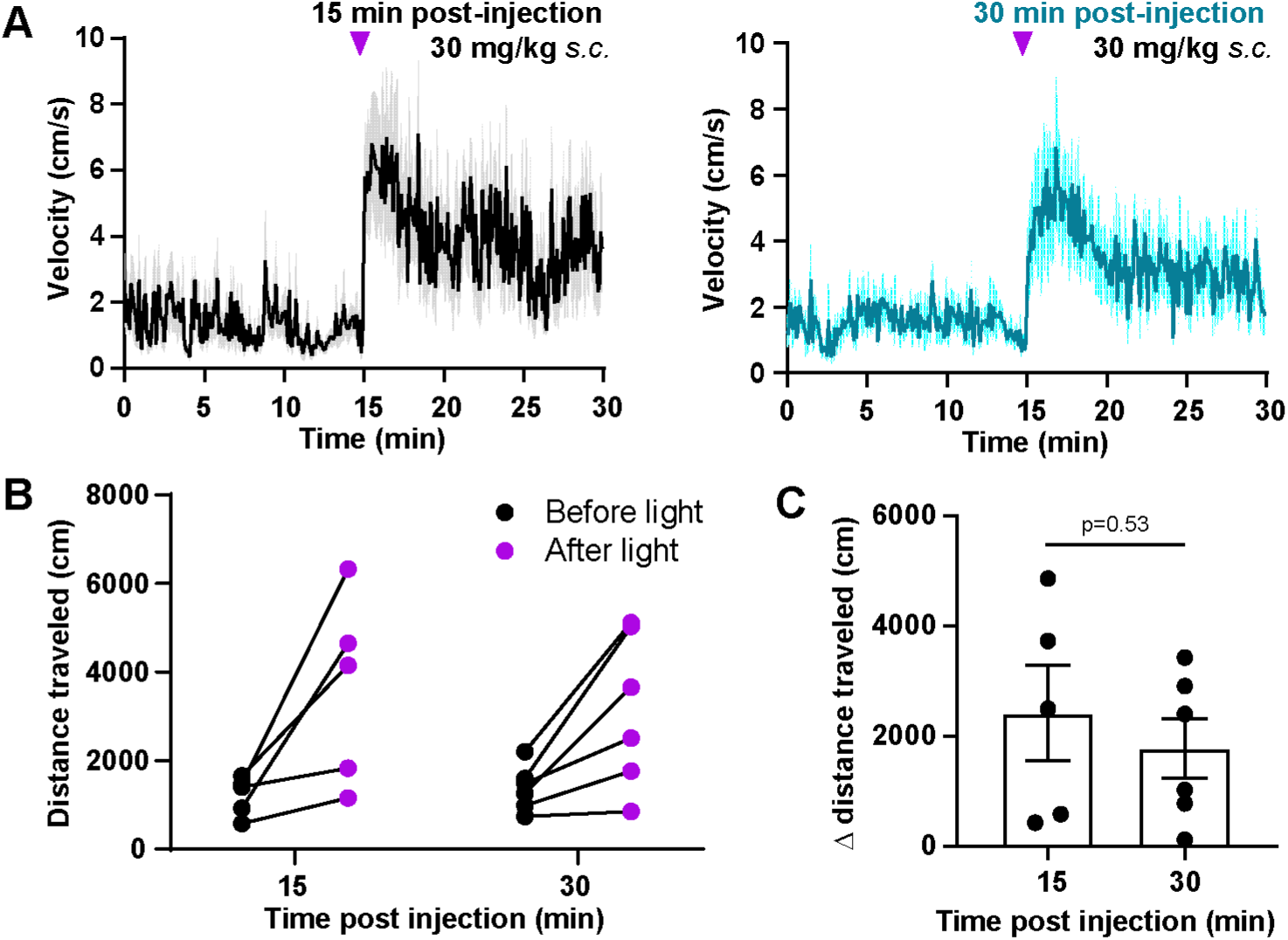
Comparison of locomotor responses to uncaging at 15 and 30 min after PhOX administration. **(A)** Average plots of velocity over time in the open field using 10 x 200 ms flashes of 375 nm light (2 Hz, 70 mW) either 15 min or 30 min after PhOX (15 mg/kg *i.v.*) administration (15 min: n = 5 mice, 30 min: n=6 mice). **(B)** Plot of the total distance traveled in the 15 minutes before and after photoactivation. **(C)** Summary plot of the change in total distance traveled in the 15 minutes before and after light delivery (Unpaired two-tailed t-test, 15 min: n=5 mice, 30 min: n=6 mice). All data are plotted as mean ± SEM.

**Figure S8.**
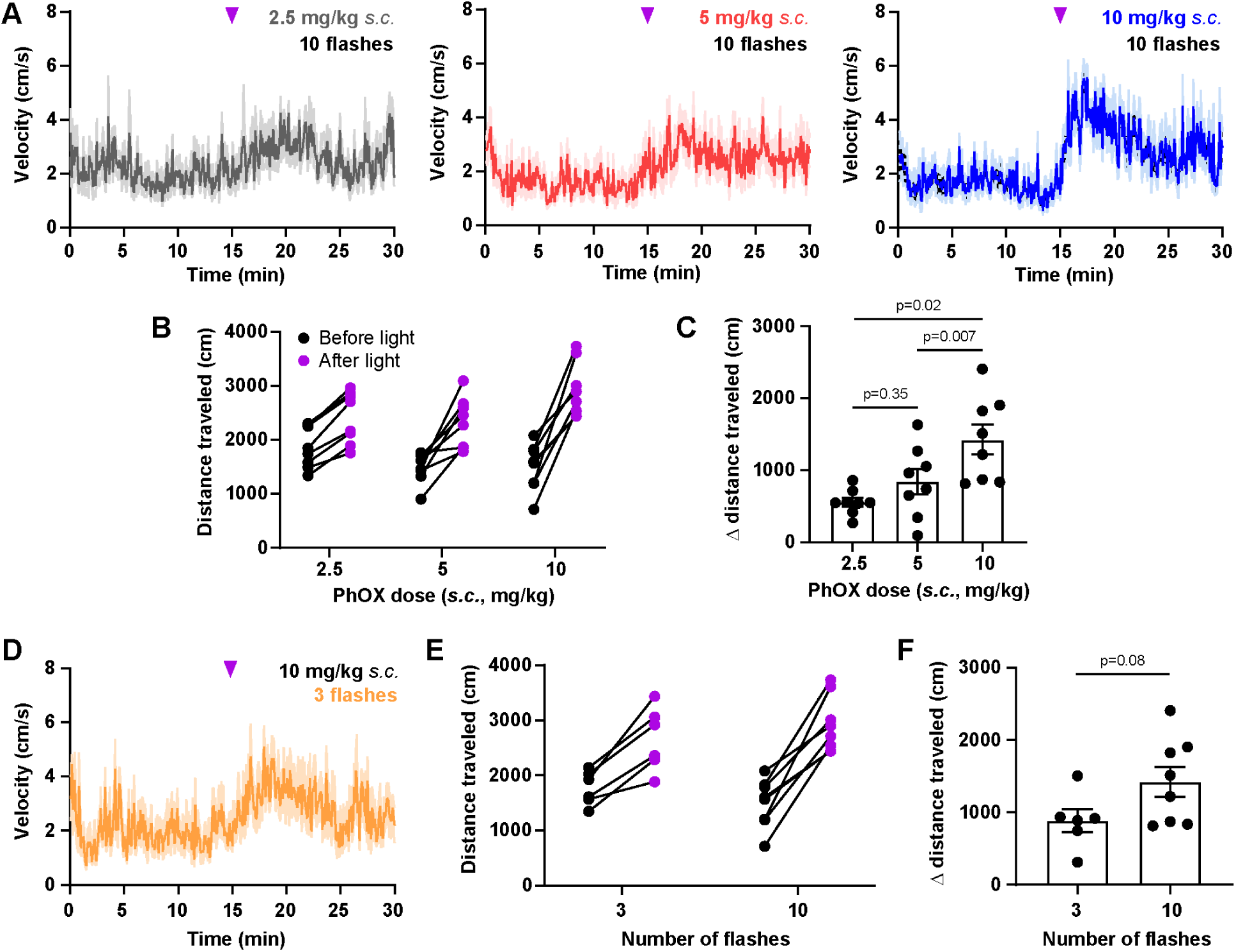
Comparison of locomotor responses to mild PhOX photoactivation conditions using subcutaneous administration. **(A)** Average plots of velocity over time in the open field using 10 x 200 ms flashes of 375 nm light (2 Hz, 70 mW) at the indicated doses of PhOX (n = 8 mice). **(B)** Plot of the total distance traveled in the 15 minutes before and after photoactivation. **(C)** Summary plot of the change in total distance traveled in the 15 minutes before and after light delivery (n=8 mice, Repeated measures one-way ANOVA, F(1.41,8.84)=10.32, p=0.006, Tukey’s multiple comparisons test). **(D)** Average plots of velocity over time in the open field using 3 x 200 ms flashes of 375 nm light (2 Hz, 70 mW) at 10 mg/kg PhOX (n = 6 mice). **(E)** Plot of the total distance traveled in the 15 minutes before and after photoactivation (10 flash data are the same as in A-C). **(F)** Summary plot of the change in total distance traveled in the 15 minutes before and after light delivery (Unpaired two-tailed t test, 3 flashes: n=6 mice, 10 flashes: n=8 mice). Although 10 flashes produced a larger average locomotor response than 3 flashes, the difference was not statistically significant. All data are plotted as mean ± SEM.

**Figure S9.**
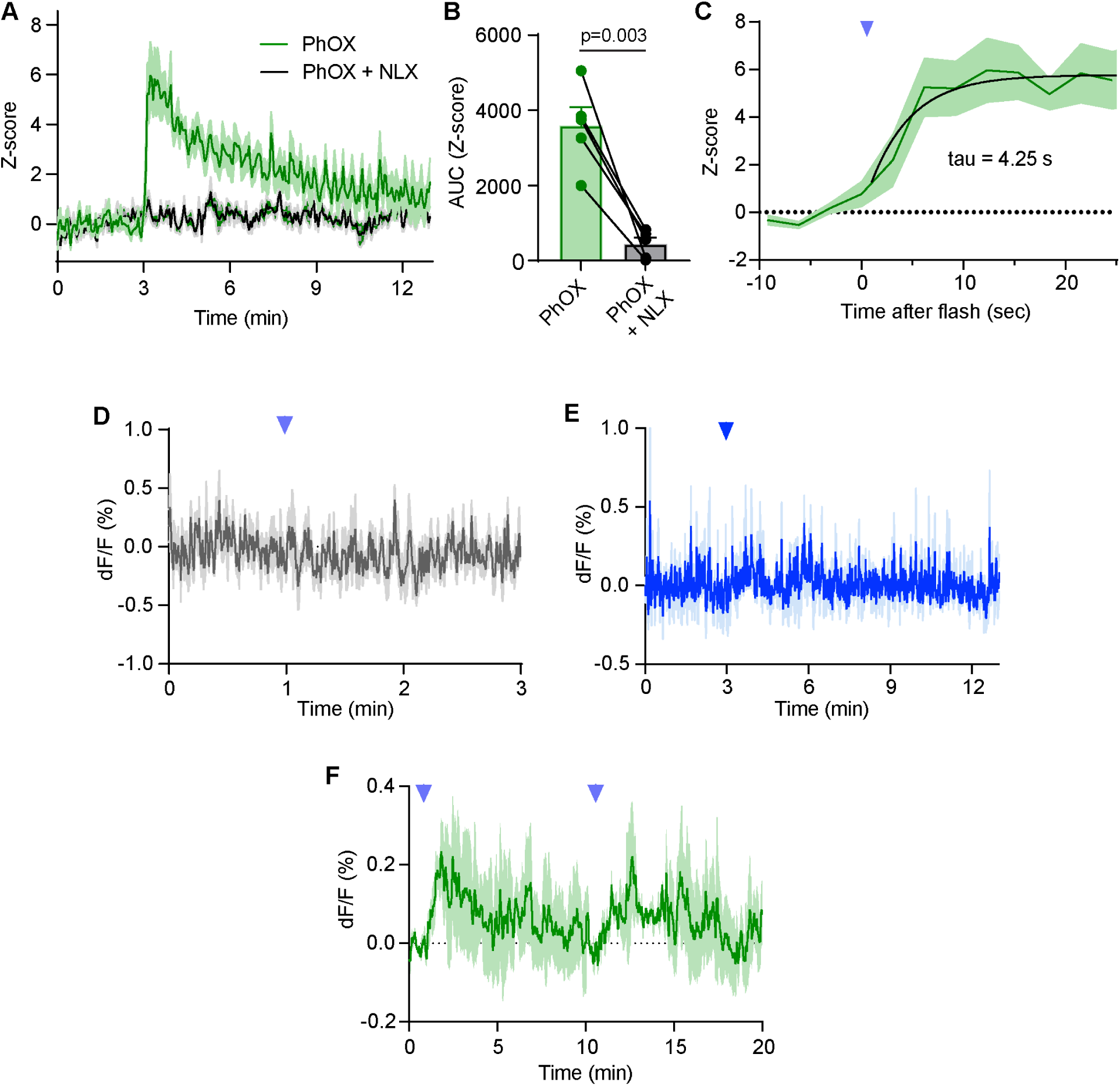
Further characterization of NAc-mSh dopamine release evoked by PhOX photoactivation in the VTA. **(A)** Average dLight1.3b fluorescence (Z-score) in the NAc-mSh in response to PhOX uncaging with a single 375 nm light flash in the ipsilateral VTA (200 ms, 50 mW) after systemic injection of either PhOX (15 mg/kg *i.v.*) or PhOX + NLX (15 mg/kg & 10 mg/kg, *i.v.*, n=5 mice). Same data shown in Figure 7C (n=5 mice). **(B)** Summary plot of the uncaging-evoked dlight1.3b fluorescence changes (Z-score) shown in (A) (AUC=area under the curve, n=5 mice, paired Student’s t-test). **(C)** Zoom-in of the rising phase of the dLight1.3b signal shown in Supporting Figure 10A (n=5 mice). **(D)** Average dLight1.3b fluorescence in the NAc-mSh in response to a single 375 nm light flash in the ipsilateral VTA (200 ms, 50 mW) in the absence of PhOX (n=4 mice). **(E)** Average dLight1.3b fluorescence in the NAc-mSh in response to 473 nm light flashes (10 x 200 ms, 50 mW, 1 Hz) in the ipsilateral VTA (200 ms, 50 mW) after systemic injection of PhOX (15 mg/kg *i.v.*) (n=2 mice). **(F)** Average dLight1.3b fluorescence in the NAc-mSh in response to repeated PhOX uncaging with two single 375 nm light flashes in the ipsilateral VTA (200 ms, 20 mW) after systemic injection of PhOX (10 mg/kg *s.c.*) (n=3 mice). All data are plotted as mean ± SEM.

**Figure S10.**
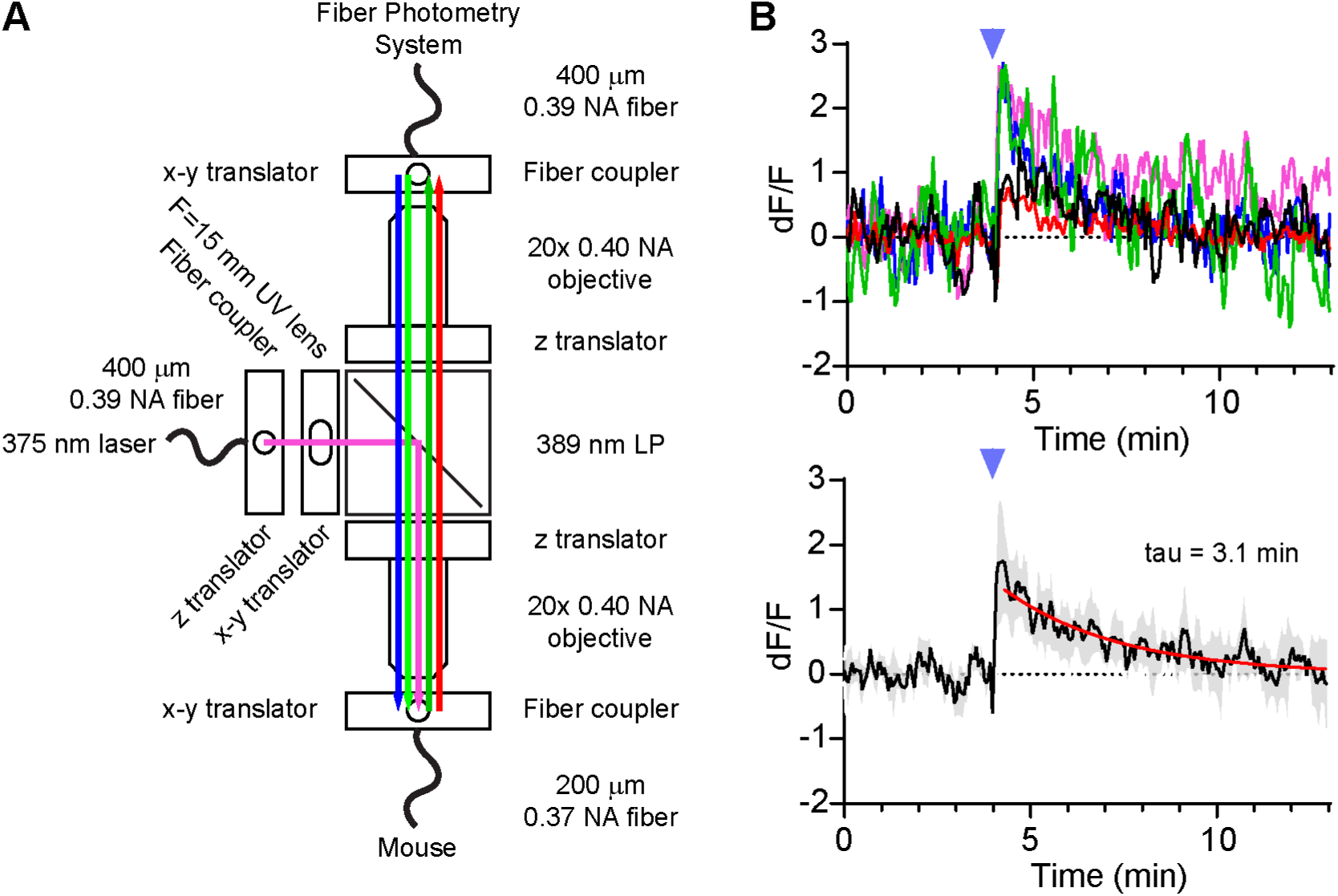
Details of the optical relay used combining UV uncaging with fiber photometry. **(A)** Schematic depicting the fiber photometry relay system used to integrate 375 nm laser light into the fiber photometry output pathway. **(B)** Fluorescence artifact produced by UV excitation of jGCaMP8s with a single light flash in the absence of PhOX (200 ms, 20 mW). Average fluorescence responses to 3 UV flashes, applied every 10 min, in individual mice (top) and average fluorescence response of all mice combined (bottom, single exponential least-squares curve fit, n=5 mice). The average artifacts from individual mice (top) were removed from the data obtained in the presence of PhOX to generate the artifact-corrected data shown in Figure 7K. All data are plotted as mean ± SEM.

**Figure S11.**
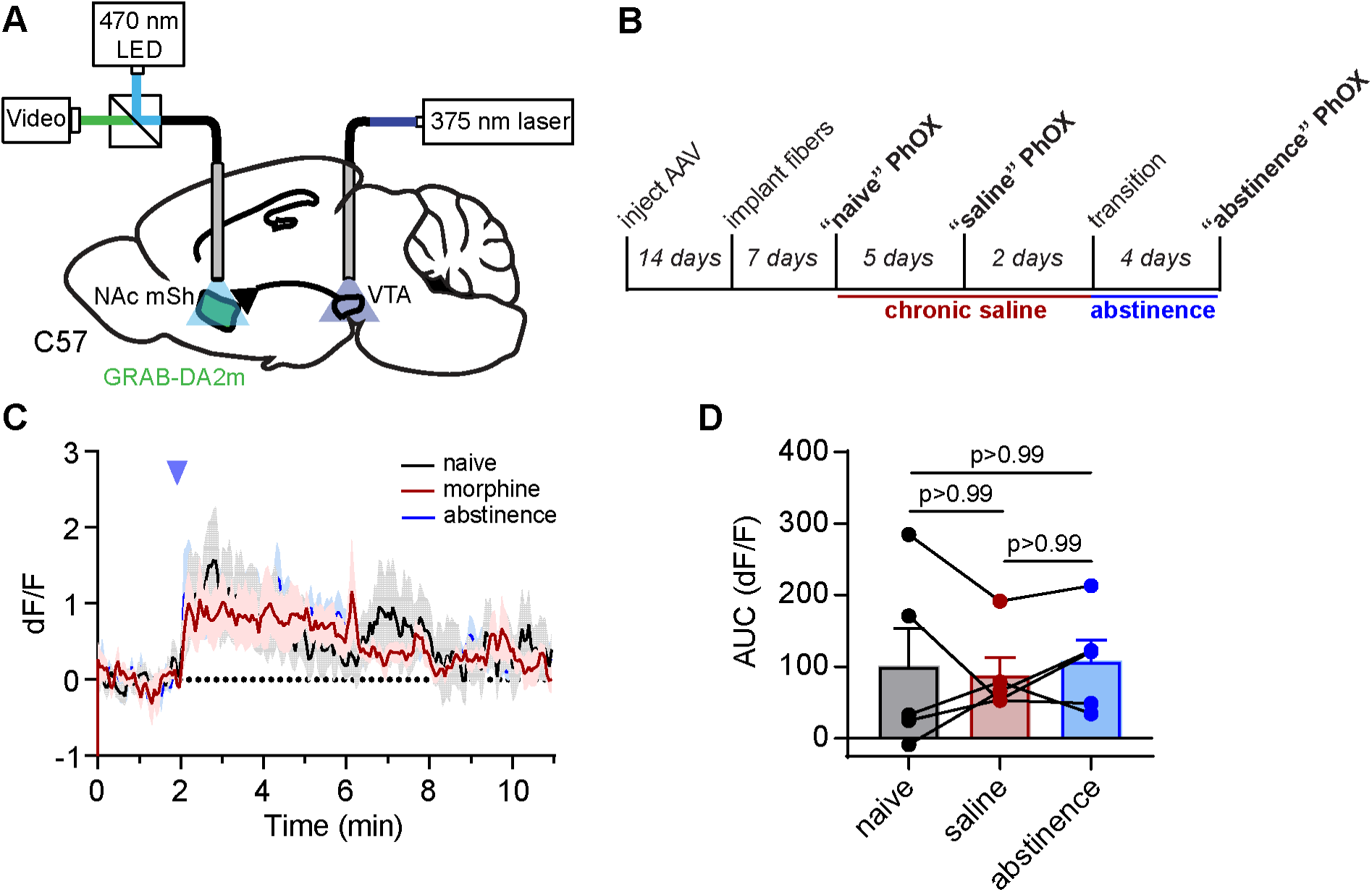
Chronic saline administration does not affect VTA PhOX uncaging-evoked dopamine release in the NAc-mSh. **(A)** Schematic indicating the implantation of a 375 nm laser-coupled fiber in the VTA and a fiber photometry-coupled fiber in the NAc-mSh to detect extracellular dopamine with dLight1.3b in response to PhOX uncaging. **(B)** Experimental timeline for fiber photometry recordings of VTA PhOX uncaging-evoked dopamine release in the NAc-mSh during chronic saline administration and abstinence. **(C)** Average GRABDA-2m fluorescence in the NAc-mSh in response to PhOX uncaging 375 nm light flashes in the VTA (10 x 200 ms, 1 Hz, 20 mW) before, during, and after chronic saline administration. **(D)** Summary plot of the uncaging-evoked GRABDA-2m fluorescence changes shown in (C) (n=5 mice, Friedman test, p=0.95, Dunn’s multiple comparisons test). All data are plotted as mean ± SEM.

## References

Acosta-Ruiz, A. et al. (2020) ‘Branched Photoswitchable Tethered Ligands Enable Ultra-efficient Optical Control and Detection of G Protein-Coupled Receptors In Vivo’, Neuron. Cell Press, 105(3), pp. 446–463.e13. doi: 10.1016/j.neuron.2019.10.036.

Banghart, M. R. et al. (2013) ‘Caged naloxone reveals opioid signaling deactivation kinetics.’, Molecular pharmacology, 84(5), pp. 687–95. doi: 10.1124/mol.113.088096.

Banghart, M. R., He, X. J. and Sabatini, B. L. (2018) ‘A Caged Enkephalin Optimized for Simultaneously Probing Mu and Delta Opioid Receptors.’, ACS chemical neuroscience, 9(4), pp. 684–690. doi: 10.1021/acschemneuro.7b00485.

Banghart, M. R. and Sabatini, B. L. (2012) ‘Photoactivatable neuropeptides for spatiotemporally precise delivery of opioids in neural tissue.’, Neuron, 73(2), pp. 249–59. doi: 10.1016/j.neuron.2011.11.016.

Bonaventura, J. et al. (2021) ‘Pharmacological and behavioral divergence of ketamine enantiomers:implications for abuse liability’, *Molecular Psychiatry*. Mol Psychiatry, 26(11), pp. 6704–6722. doi: 10.1038/s41380-021-01093-2.

Borg, P. J. and Taylor, D. A. (1997) ‘Involvement of μ- and δ-opioid receptors in the effects of systemic and locally perfused morphine on extracellular levels of dopamine, DOPAC and HVA in the nucleus accumbens of the halothane-anaesthetized rat’, *Naunyn-Schmiedeberg’s Archives of Pharmacology*. Naunyn Schmiedebergs Arch Pharmacol, 355(5), pp. 582–588. doi: 10.1007/PL00004987.

Britt, J. P. and McGehee, D. S. (2008) ‘Presynaptic opioid and nicotinic receptor modulation of dopamine overflow in the nucleus accumbens’, Journal of Neuroscience. Society for Neuroscience, 28(7), pp. 1672–1681. doi: 10.1523/JNEUROSCI.4275-07.2008.

Broichhagen, J., Frank, J. A. and Trauner, D. (2015) ‘A Roadmap to Success in Photopharmacology’, Accounts of Chemical Research. American Chemical Society, 48(7), pp. 1947–1960. doi: 10.1021/acs.accounts.5b00129.

Bruno, C. A. et al. (2021) ‘pMAT: An open-source software suite for the analysis of fiber photometry data’, *Pharmacology Biochemistry and Behavior*. Pharmacol Biochem Behav, 201. doi: 10.1016/j.pbb.2020.173093.

Cai, N. S. et al. (2019) ‘Opioid-galanin receptor heteromers mediate the dopaminergic effects of opioids’, *Journal of Clinical Investigation*. J Clin Invest, 129(7), pp. 2730–2744. doi: 10.1172/JCI126912.

Di Chiara, G. and Imperato, A. (1988a) ‘Drugs abused by humans preferentially increase synaptic dopamine concentrations in the mesolimbic system of freely moving rats’, *Proceedings of the National Academy of Sciences of the United States of America*. Proc Natl Acad Sci U S A, 85(14), pp. 5274–5278. doi: 10.1073/pnas.85.14.5274.

Di Chiara, G. and Imperato, A. (1988b) ‘Opposite effects of mu and kappa opiate agonists on dopamine release in the nucleus accumbens and in the dorsal caudate of freely moving rats’, Journal of Pharmacology and Experimental Therapeutics, 244(3), pp. 1067–1080.

Christie, M. J. (2008) ‘Cellular neuroadaptations to chronic opioids: Tolerance, withdrawal and addiction’, *British Journal of Pharmacology*. Br J Pharmacol, pp. 384–396. doi: 10.1038/bjp.2008.100.

Corre, J. et al. (2018) ‘Dopamine neurons projecting to medial shell of the nucleus accumbens drive heroin reinforcement’, eLife, 7. doi: 10.7554/eLife.39945.

Daglish, M. R. C. et al. (2008) ‘Brain dopamine response in human opioid addiction’, *British Journal of Psychiatry*. Br J Psychiatry, 193(1), pp. 65–72. doi: 10.1192/bjp.bp.107.041228.

Donthamsetti, P. et al. (2021) ‘Cell specific photoswitchable agonist for reversible control of endogenous dopamine receptors’, *Nature Communications*. Nat Commun, 12(1). doi: 10.1038/s41467-021-25003-w.

Donthamsetti, P. C. et al. (2019) ‘Genetically Targeted Optical Control of an Endogenous G Protein-Coupled Receptor’, Journal of the American Chemical Society, 141(29), pp. 11522– 11530. doi: 10.1021/jacs.9b02895.

Ellis-Davies, G. C. R. (2007) ‘Caged compounds: photorelease technology for control of cellular chemistry and physiology’, Nature Methods, 4(8), pp. 619–628. doi: 10.1038/nmeth1072.

Font, J. et al. (2017) ‘Optical control of pain in vivo with a photoactive mGlu5 receptor negative allosteric modulator’, eLife. eLife Sciences Publications Ltd, 6. doi: 10.7554/eLife.23545.

Franklin, K. B. J. (1989) ‘Analgesia and the neural substrate of reward’, *Neuroscience and Biobehavioral Reviews*. Neurosci Biobehav Rev, 13(2–3), pp. 149–154. doi: 10.1016/S0149-7634(89)80024-7.

Glickfeld, L. L., Atallah, B. V. and Scanziani, M. (2008) ‘Complementary modulation of somatic inhibition by opioids and cannabinoids’, Journal of Neuroscience, 28(8), pp. 1824–1832. doi: 10.1523/JNEUROSCI.4700-07.2008.

Gomila, A. M. J. et al. (2020) ‘Photocontrol of Endogenous Glycine Receptors In Vivo’, *Cell Chemical Biology*. Cell Chem Biol, 27(11), pp. 1425–1433.e7. doi: 10.1016/j.chembiol.2020.08.005.

Gorostiza, P. and Isacoff, E. Y. (2008) ‘Optical switches for remote and noninvasive control of cell signaling’, *Science*. Science, pp. 395–399. doi: 10.1126/science.1166022.

Gysling, K. and Wang, R. Y. (1983) ‘Morphine-induced activation of A10 dopamine neurons in the rat’, *Brain Research*. Brain Res, 277(1), pp. 119–127. doi: 10.1016/0006-8993(83)90913-7.

Handal, M. et al. (2002) ‘Pharmacokinetic differences of morphine and morphine-glucuronides are reflected in locomotor activity’, Pharmacology Biochemistry and Behavior. Elsevier, 73(4), pp. 883–892. doi: 10.1016/S0091-3057(02)00925-5.

He, X. J. et al. (2021) ‘Convergent, functionally independent signaling by mu and delta opioid receptors in hippocampal parvalbumin interneurons’, eLife. eLife Sciences Publications Ltd, 10. doi: 10.7554/eLife.69746.

Hüll, K., Morstein, J. and Trauner, D. (2018) ‘In Vivo Photopharmacology’, Chemical Reviews. American Chemical Society, pp. 10710–10747. doi: 10.1021/acs.chemrev.8b00037.

Jalabert, M. et al. (2011) ‘Neuronal circuits underlying acute morphine action on dopamine neurons’, *Proceedings of the National Academy of Sciences of the United States of America*. Proc Natl Acad Sci U S A, 108(39), pp. 16446–16450. doi: 10.1073/pnas.1105418108.

Johnson, S. W. and North, R. A. (1992) ‘Opioids excite dopamine neurons by hyperpolarization of local interneurons’, *Journal of Neuroscience*. J Neurosci, 12(2), pp. 483–488. doi: 10.1523/jneurosci.12-02-00483.1992.

Joyce, E. M. and Iversen, S. D. (1979) ‘The effect of morphine applied locally to mesencephalic dopamine cell bodies on spontaneous motor activity in the rat’, *Neuroscience Letters*. Neurosci Lett, 14(2–3), pp. 207–212. doi: 10.1016/0304-3940(79)96149-4.

Leone, P., Pocock, D. and Wise, R. A. (1991) ‘Morphine-dopamine interaction: Ventral tegmental morphine increases nucleus accumbens dopamine release’, *Pharmacology, Biochemistry and Behavior*. Pharmacol Biochem Behav, 39(2), pp. 469–472. doi: 10.1016/0091-3057(91)90210-S.

Levitz, J. et al. (2016) ‘A toolkit for orthogonal and in vivo optical manipulation of ionotropic glutamate receptors’, Frontiers in Molecular Neuroscience. Frontiers Media S.A., 9(FEB), p. 2. doi: 10.3389/fnmol.2016.00002.

Lima, S. Q. and Miesenböck, G. (2005) ‘Remote control of behavior through genetically targeted photostimulation of neurons’, *Cell*. Cell, 121(1), pp. 141–152. doi: 10.1016/j.cell.2005.02.004.

Lin, W. C. et al. (2015) ‘A Comprehensive Optogenetic Pharmacology Toolkit for In Vivo Control of GABAA Receptors and Synaptic Inhibition’, *Neuron*. Neuron, 88(5), pp. 879–891. doi: 10.1016/j.neuron.2015.10.026.

López-Cano, M. et al. (2021) ‘Remote local photoactivation of morphine produces analgesia without opioid-related adverse effects’, *British Journal of Pharmacology*. Br J Pharmacol. doi: 10.1111/bph.15645.

Ma, Y. et al. (2005) ‘A three-dimensional digital atlas database of the adult C57BL/6J mouse brain by magnetic resonance microscopy’, *Neuroscience*. Neuroscience, 135(4), pp. 1203–1215. doi: 10.1016/j.neuroscience.2005.07.014.

Manning, B. H., Morgan, M. J. and Franklin, K. B. J. (1994) ‘Morphine analgesia in the formalin test: Evidence for forebrain and midbrain sites of action’, *Neuroscience*. Neuroscience, 63(1), pp. 289–294. doi: 10.1016/0306-4522(94)90023-X.

Matsui, A. et al. (2014) ‘Separate GABA afferents to dopamine neurons mediate acute action of opioids, development of tolerance, and expression of withdrawal’, *Neuron*. Neuron, 82(6), pp. 1346–1356. doi: 10.1016/j.neuron.2014.04.030.

Matsui, A. and Williams, J. T. (2011) ‘Opioid-sensitive GABA inputs from rostromedial tegmental nucleus synapse onto midbrain dopamine neurons’, Journal of Neuroscience, 31(48), pp. 17729–17735. doi: 10.1523/JNEUROSCI.4570-11.2011.

Mourot, A. et al. (2011) ‘Tuning photochromic ion channel blockers’, ACS Chemical Neuroscience. American Chemical Society, 2(9), pp. 536–543. doi: 10.1021/cn200037p.

Olmstead, M. C. and Franklin, K. B. J. (1997) ‘The development of a conditioned place preference to morphine: effects of microinjections into various CNS sites’, *Behavioral neuroscience*. Behav Neurosci, 111(6), pp. 1324–1334. doi: 10.1037//0735-7044.111.6.1324.

Owen, S. F., Liu, M. H. and Kreitzer, A. C. (2019) ‘Thermal constraints on in vivo optogenetic manipulations’, Nature Neuroscience. Nature Publishing Group, 22(7), pp. 1061–1065. doi: 10.1038/s41593-019-0422-3.

Packer, A. M. et al. (2015) ‘Simultaneous all-optical manipulation and recording of neural circuit activity with cellular resolution in vivo’, *Nature Methods*. Nat Methods, 12(2), pp. 140–146. doi: 10.1038/nmeth.3217.

Pentney, R. J. W. and Gratton, A. (1991) ‘Effects of local delta and mu opioid receptor activation on basal and stimulated dopamine release in striatum and nucleus accumbens of rat: An in vivo electrochemical study’, *Neuroscience*. Neuroscience, 45(1), pp. 95–102. doi: 10.1016/0306-4522(91)90106-X.

Rashid, H. et al. (2003) ‘Novel expression of vanilloid receptor 1 on capsaicin-insensitive fibers accounts for the analgesic effect of capsaicin cream in neuropathic pain’, *Journal of Pharmacology and Experimental Therapeutics*. J Pharmacol Exp Ther, 304(3), pp. 940–948. doi: 10.1124/jpet.102.046250.

Ren, W., Ji, A. and Ai, H. W. (2015) ‘Light activation of protein splicing with a photocaged fast intein’, *Journal of the American Chemical Society*. J Am Chem Soc, 137(6), pp. 2155–2158. doi: 10.1021/ja508597d.

Schaal, J. et al. (2009) ‘A novel photorearrangement of (coumarin-4-yl)methyl phenyl ethers’, Journal of Photochemistry and Photobiology A: Chemistry. Elsevier, 208(2–3), pp. 171–179. doi: 10.1016/j.jphotochem.2009.09.012.

Shields, B. C. et al. (2017) ‘Deconstructing behavioral neuropharmacology with cellular specificity’, Science, 356(6333), p. eaaj2161. doi: 10.1126/science.aaj2161.

Spagnolo, P. A. et al. (2019) ‘Striatal Dopamine Release in Response to Morphine: A [11C]Raclopride Positron Emission Tomography Study in Healthy Men’, Biological Psychiatry. NIH Public Access, 86(5), pp. 356–364. doi: 10.1016/j.biopsych.2019.03.965.

Spanagel, R., Herz, A. and Shippenberg, T. S. (1990) ‘The Effects of Opioid Peptides on Dopamine Release in the Nucleus Accumbens: An In Vivo Microdialysis Study’, *Journal of Neurochemistry*. J Neurochem, 55(5), pp. 1734–1740. doi: 10.1111/j.1471-4159.1990.tb04963.x.

Sun, F. et al. (2020) ‘Next-generation GRAB sensors for monitoring dopaminergic activity in vivo’, Nature Methods. Nature Research, 17(11), pp. 1156–1166. doi: 10.1038/s41592-020-00981-9.

Tanowitz, M. and Von Zastrow, M. (2003) ‘A Novel Endocytic Recycling Signal That Distinguishes the Membrane Trafficking of Naturally Occurring Opioid Receptors’, Journal of Biological Chemistry. Elsevier, 278(46), pp. 45978–45986. doi: 10.1074/jbc.M304504200.

Taylor, A. M. W. et al. (2016) ‘Neuroimmune Regulation of GABAergic Neurons Within the Ventral Tegmental Area during Withdrawal from Chronic Morphine’, *Neuropsychopharmacology*. Neuropsychopharmacology, 41(4), pp. 949–959. doi: 10.1038/npp.2015.221.

Tobias, J. M. et al. (2021) ‘Genetically-targeted photorelease of endocannabinoids enables optical control of GPR55 in pancreatic β-cells’, *Chemical Science*. Chem Sci, 12(40), pp. 13506– 13512. doi: 10.1039/d1sc02527a.

Velema, W. A., Szymanski, W. and Feringa, B. L. (2014) ‘Photopharmacology: Beyond proof of principle’, Journal of the American Chemical Society. American Chemical Society, pp. 2178– 2191. doi: 10.1021/ja413063e.

Wang, J. B. et al. (2018) ‘Noninvasive Ultrasonic Drug Uncaging Maps Whole-Brain Functional Networks’, *Neuron*. Neuron, 100(3), pp. 728–738.e7. doi: 10.1016/j.neuron.2018.10.042.

Watson, B. J. et al. (2014) ‘Investigating expectation and reward in human opioid addiction with [11C]raclopride PET’, *Addiction Biology*. Addict Biol, 19(6), pp. 1032–1040. doi: 10.1111/adb.12073.

Williams, J. T. et al. (2013) ‘Regulation of μ-opioid receptors: Desensitization, phosphorylation, internalization, and tolerance’, *Pharmacological Reviews*. Pharmacol Rev, pp. 223–254. doi: 10.1124/pr.112.005942.

Williams, J. T. (2014) ‘Desensitization of Functional -Opioid Receptors Increases Agonist Off-Rate’, Molecular Pharmacology, 86(1), pp. 52–61. doi: 10.1124/mol.114.092098.

Wong, P. T. et al. (2017) ‘Control of an Unusual Photo-Claisen Rearrangement in Coumarin Caged Tamoxifen through an Extended Spacer’, ACS Chemical Biology. American Chemical Society, 12(4), pp. 1001–1010. doi: 10.1021/acschembio.6b00999.

Wootton, J. F. and Trentham, D. R. (1989) ‘“Caged” compounds to probe the dynamics of cellular processes: synthesis and properties of some novel photosensitive P-2-nitrobenzyl esters of nucleotides’, in Nielsen, P. E. (ed.) Photochemical Probes In Biochemistry. NATO ASI S. Kluwer Academic, The Netherlands, pp. 277–296.

Yaksh, T. L., Yeung, J. C. and Rudy, T. A. (1976) ‘Systematic examination in the rat of brain sites sensitive to the direct application of morphine: observation of differential effects within the periaqueductal gray.’, Brain research, 114(1), pp. 83–103. Available at: http://www.ncbi.nlm.nih.gov/pubmed/963546 (Accessed: 28 November 2018).

Zhang, Y. et al. (2021) ‘Fast and sensitive GCaMP calcium indicators for imaging neural populations’, *bioRxiv*. Cold Spring Harbor Laboratory, p. 2021.11.08.467793. doi: 10.1101/2021.11.08.467793.

